# Modular prophage interactions driven by capsule serotype select for capsule loss under phage predation

**DOI:** 10.1101/2019.12.17.878363

**Authors:** Jorge A. Moura de Sousa, Amandine Buffet, Matthieu Haudiquet, Eduardo P.C. Rocha, Olaya Rendueles

## Abstract

*Klebsiella* species are able to colonize a wide range of environments and include worrisome nosocomial pathogens. Here, we sought to determine the abundance and infectivity of prophages of *Klebsiella* to understand how the interactions between induced prophages and bacteria affect population dynamics and evolution. We identified many prophages in the species, placing these taxa among the top 5% of the most polylysogenic bacteria. We selected 35 representative strains of the *Klebsiella pneumoniae* species complex to establish a network of induced phage-bacteria interactions. This revealed that many prophages are able to enter the lytic cycle, and subsequently kill or lysogenize closely-related *Klebsiella* strains. Although 60% of the tested strains could produce phages that infect at least one other strain, the interaction network of all pairwise cross-infections is very sparse and mostly organized in modules corresponding to the strains’ capsule serotypes. Accordingly, capsule mutants remain uninfected showing that the capsule is a key factor for successful infections. Surprisingly, experiments in which bacteria are predated by their own prophages result in accelerated loss of the capsule. Our results show that phage infectiousness defines interaction modules between small subsets of phages and bacteria in function of capsule serotype. This limits the role of prophages as competitive weapons because they can infect very few strains of the species complex. This should also restrict phage-driven gene flow across the species. Finally, the accelerated loss of the capsule in bacteria being predated by their own phages, suggests that phages drive serotype switch in nature.

## INTRODUCTION

Phages are one of the most abundant entities on Earth. They are found in multiple environments, typically along with their host bacteria, including in the human microbiome. Many recent studies focused on virulent phages, which follow exclusively a lytic cycle. In contrast, temperate phages, which can either follow a lytic cycle, or integrate into the host genome and produce a lysogen, have been comparatively less studied. Integrated phages, hereafter referred to as prophages, replicate vertically with the host and are typically able to protect them from infections by similar phages, the so-called resistance to superinfection (Canchaya et al 2003, Lu and Henning 1994, Susskind et al 1974). Most prophage genes are silent and have little impact in bacterial fitness as long as there is no induction of the lytic cycle (Canchaya et al 2003). If the prophage remains in the genome for a very long period of time it may be inactivated by mutations. A few studies suggest that many prophages are inactive to some extent (Asadulghani et al 2009, Matos et al 2013). Upon induction, some of them cannot excise (cryptic prophages), replicate, infect, or produce viable progeny. Prophage inactivation and further domestication may lead to the co-option of some phage functions by the bacterial host (Touchon et al 2014a). For instance, some bacteriocins result from the domestication of phage tails (Nakayama et al 2000, Winstanley et al 2009). In contrast, intact prophages can be induced (by either extrinsic or intrinsic factors) and resume a lytic cycle, producing viable viral particles.

Temperate phages affect the evolution of gene repertoires and bacterial population dynamics (Bossi et al 2003, Fortier and Sekulovic 2013, Nanda et al 2015), by two key mechanisms. First, induction of prophages by a small subset of a population produces virions that can infect susceptible bacteria and thus facilitate colonization (Bossi et al 2003, Brown et al 2006, Sousa and Rocha 2019). Second, they drive horizontal gene transfer between bacteria. Around half of the sequenced genomes contain identifiable prophages (Roux et al 2015, Touchon et al 2016), e.g. a third of *E. coli*’s pangenome is in prophages (Bobay et al 2013). The frequency of prophages is higher in bacteria with larger genomes, in pathogens, and in fast-growing bacteria (Touchon et al 2016). Lysogenization and the subsequent expression of some prophage genes may result in phenotypic changes in the host, e.g., many pathogens have virulence factors encoded in prophages (Wagner and Waldor 2002). Prophages may also facilitate horizontal transfer between bacteria by one of several mechanisms of transduction (Canchaya et al 2003, Chen et al 2018, Touchon et al 2017). Interestingly, bacterial populations can acquire adaptive genes from their competitors by killing them with induced prophages and recovering their genes by generalized transduction (Haaber et al 2016). While these mechanisms have been explored in many experimental and computational studies, the impact of temperate phages in the diversity of bacterial lysogens is still poorly understood.

Here, we assess the relevance of prophages in the biology of *Klebsiella* spp., a genus of bacteria capable of colonizing a large range of environments. The genus includes genetically diverse species of heterotrophic facultative aerobes that have been isolated from numerous environments, including the soil, sewage, water, plants, and mammals (Brisse et al 2006). *Klebsiella spp.* can cause various diseases such as urinary tract infections, acute liver abscesses, pneumonia, infectious wounds and dental infections (Lee et al 2017, Navon-Venezia et al 2017). They commonly cause severe hospital outbreaks associated with multidrug resistance (MDR), and *K. pneumoniae* is one of the six most worrisome antibiotic-resistant (ESKAPE) pathogens. The versatility of *Klebsiella spp*. is associated with a broad and diverse metabolism (Blin et al 2017), partly acquired by horizontal gene transfer (Holt et al 2015, Navon-Venezia et al 2017, Wyres et al 2019). Additionally, *Klebsiella spp*. code for an extracellular capsule that is highly variable within the species. This capsule is a high molecular weight polysaccharide made up of different repeat units of oligosaccharides. Combinations of different oligosaccharides are referred to as serotypes. In *Klebsiella pneumoniae* there are 77 serologically-defined serotypes, and numbered from K1 to K77 (Mori et al 1989) and more than 130 serotypes were identified computationally. The latter are noted from KL1 to KL130 (Pan et al 2015, Wyres et al 2016), and are usually referred as capsule locus types (or CLT). The capsule is considered a major virulence factor, required, for instance, in intestinal colonization (Favre-Bonte et al 1999). It also provides resistance to the immune response and to antibiotics (Alvarez et al 2000, Campos et al 2004, Doorduijn et al 2016). From an ecological point of view, the capsule is associated with bacteria able to colonize diverse environments (Rendueles et al 2017, Rendueles et al 2018). Its rapid diversification may thus be a major contributor to *Klebsiella’*s adaptive success, including in colonizing clinical settings.

We have recently shown that species of bacteria encoding capsular loci undergo more frequent genetic exchanges and accumulate more mobile genetic elements, including prophages (Rendueles et al 2018). This is surprising because capsules were proposed to decrease gene flow (Chewapreecha et al 2014) and some phages are known to be blocked by the capsule (Moller et al 2019, Negus et al 2013, Scholl et al 2005). However, several virulent phages of *Klebsiella* are known to have depolymerase activity in their tail fibers (Bessler et al 1973, Niemann et al 1977a, Niemann et al 1977b, Pan et al 2017, Thurow et al 1974). These depolymerases specifically digest oligosaccharidic bonds in certain capsules (Latka et al 2017) and allow phages to access the outer membrane and infect bacteria (Scholl et al 2001). Since depolymerases are specific to one or a few capsule types (Lin et al 2014, Thurow et al 1974), this implicates that some phages interact with capsules in a serotype-specific manner (Pan et al 2017, Pan et al 2019, Thurow et al 1974). Additionally, the capsule could facilitate cell infection because phages bind reversibly to it, prior to the irreversible binding to the specific cell receptor (Bertozzi Silva et al 2016).

To date, the number and role of prophages in *Klebsiella*’s population biology is not well known. *Klebsiella* are interesting models to study the role of prophages, because of the interplay between the capsule, phage infections, and also the influence of the former in *Klebsiella’s* colonization of very diverse ecological niches. In this work, we sought to characterize the abundance and distribution of *Klebsiella* temperate phages, and experimentally assess their ability to re-enter the lytic cycle and lysogenize different *Klebsiella* strains. By performing more than 1200 pairwise combinations of lysates and host strains, we aim to pinpoint the drivers of prophage distribution and elucidate some of the complex interactions that shape phage-bacteria interactions in *Klebsiella*.

## RESULTS

### Prophages are very abundant in the genomes of Klebsiella

We used PHASTER (Arndt et al 2016) to analyse 254 genomes of eight *Klebsiella* species (and two subspecies). We detected 1674 prophages, of which 55% are classified as “intact” by PHASTER and are the most likely to be complete and functional. These “intact” prophages were present in 237 out of the 254 genomes (see Methods). The remaining prophages were classed as “questionable” (20%) and “incomplete” (25%, Figure 1AB, Figure S1). The complete list of bacterial genomes and prophages is available in Dataset S1. Most of the genomes were poly-lysogenic, encoding more than one prophage (Figure S1). However, the number of prophages varied markedly across genomes, ranging from one to 16, with a median of 6 per genome. Additionally, the total number of prophages in *Klebsiella* spp. varied significantly across species (Kruskal-Wallis, *df* = 6, *P* < 0.001, Figure 1B). More specifically, the average number of “intact” prophages varied between eight for *K. oxytoca* and less than one for *K. quasipneumoniae subsp. quasipneumoniae*. *Klebsiella pneumoniae*, the most represented species of our dataset (∼77% of the genomes), has an average of nine prophages per genome, of which four are classified as “intact” (Figure 1B). As expected, both the number of prophages per genome and the average number of prophages per species are correlated positively with genome size (Spearman’s rho= 0.49, P<0.001, Spearman’s rho= 0.76, P=0.01, respectively) (Figure S2AB). When compared with the one hundred most sequenced bacterial species, the number of prophages in *Klebsiella* is very high. It ranks *K. pneumoniae* within the 5^th^ percentile of the most prophage-rich species, comparable to *E. coli* and *Yersinia enterocolitica* (Figure S3). This shows that prophages are a sizeable fraction of the genomes of *Klebsiella* and may have an important impact in its biology.

**Figure 1.**
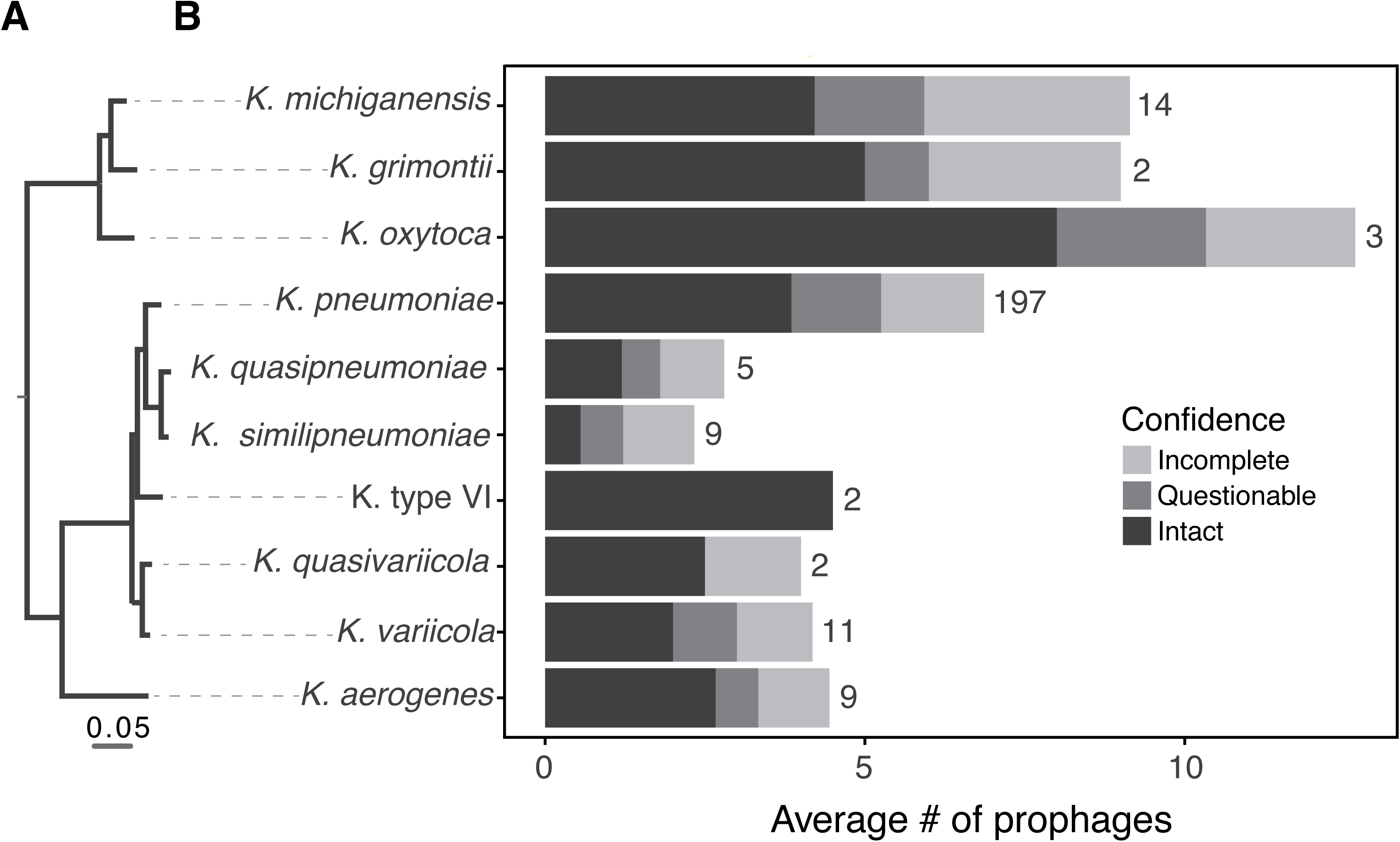
Prophage distribution in *Klebsiella* genus. **A.** Rooted phylogenetic tree of *Klebsiella* species used in this study based on the core genes. **B.** Average number of prophages per genome. PHASTER prediction for completeness is indicated. Numbers represent the total number of genomes analysed of each species.

Our experience is that prophages classed as “questionable” and “incomplete” often lack essential phage functions. Hence, all the remaining analyses were performed on “intact” prophages, unless indicated otherwise. These elements have 5% lower GC content than the rest of the chromosome (Wilcoxon test, P< 0.001, Figure S2B), as typically observed for horizontally transferred genetic elements (Daubin et al 2003, Rocha and Danchin 2002). Their length varies from 13 to 137 kb, for an average of 46kb. Since temperate dsDNA phages of enterobacteria are usually larger than 30kb (Bobay et al 2014), this suggests that a small fraction of the prophages might be incomplete (Figure S2C). Among the “intact” prophages detected in the 35 strains analysed from our laboratory collection (Dataset S1 and Figure S1) and isolated from different environments and representative of the genetic diversity of the *Klebsiella pneumoniae* species complex (Blin et al 2017), we chose eleven to characterize in detail in terms of genetic architecture (Figure 2). A manual and computational search for recombination sites (*att*) in these eleven prophages (See Methods, Figure 2) showed that some might be larger than predicted by PHASTER. To verify the integrity of these phages, we searched for genes encoding structural functions - head, baseplate and tail – and found them in the eleven prophages. Two of these prophages (#62(2) and #54 (1)) have a protein of *ca.* 4200 aa (Figure 2), homologous to the tail component (gp21) of the linear phage-plasmid phiKO2 of *Klebsiella oxytoca* (Casjens et al 2004). The prophages had integrases and were flanked by recombination sites, suggesting that they could still be able to excise themselves from the chromosome (Figure 2). Nevertheless, some prophages encoded a small number of insertion sequences (IS), which accumulate under lysogeny and degrade prophages (Leclercq and Cordaux 2011).

**Figure 2.**
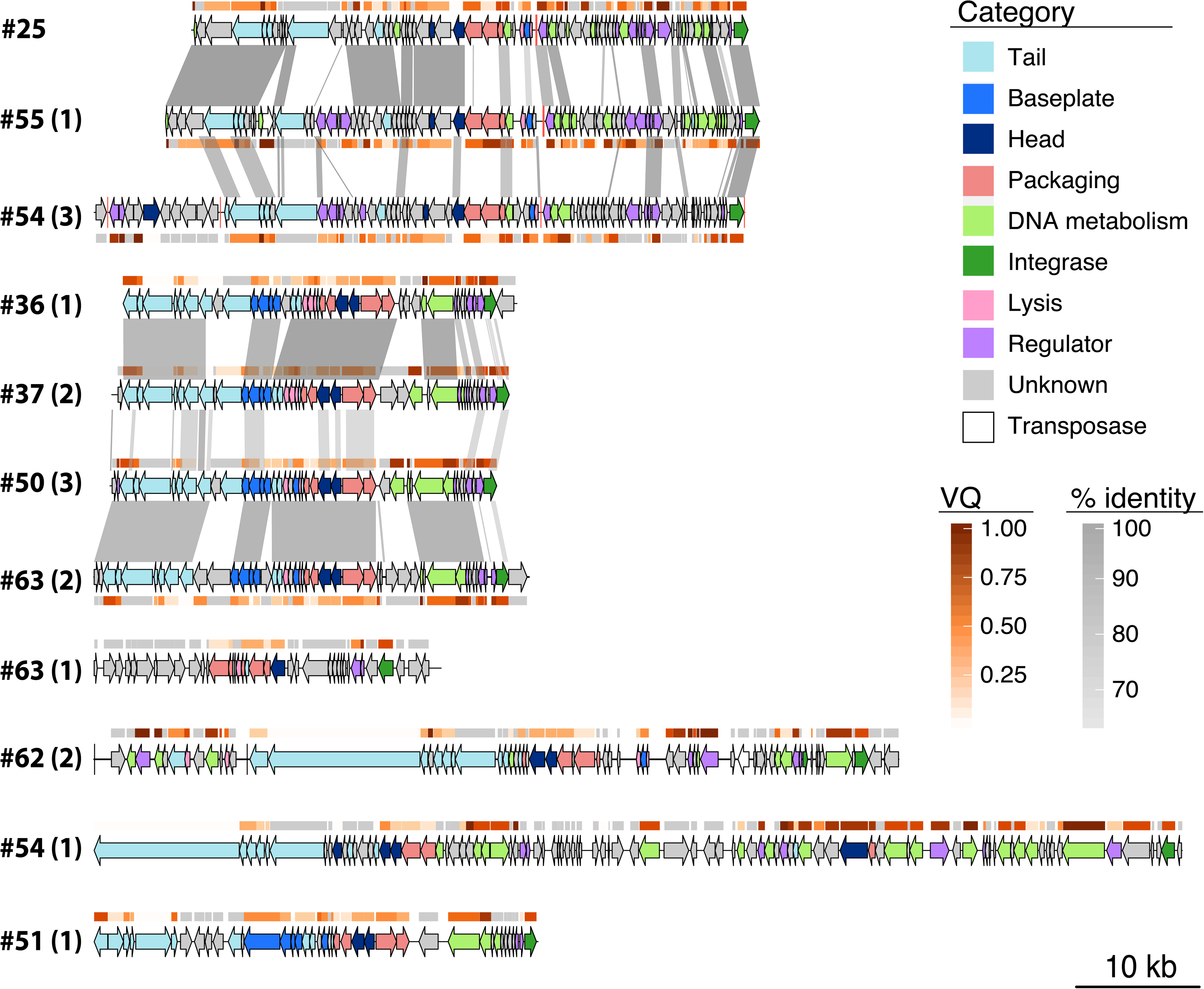
Genomic organization of nine of the prophages in this study. Numbers correspond to the host genomes, as displayed in Figure 3. The number in parenthesis identifies the prophage in the genome. All prophages are classified as “intact” except #63 (1) (“questionable”). Genome boundaries correspond to attL/R sites. Arrows represent predicted ORFs and are oriented according to transcriptional direction. Colors indicate assigned functional categories, tRNAs are represented as red lines and the sequences are oriented based on the putative integrase localization. Local *blastn* alignments (option *dc-megablast*) are displayed between pairs of related prophages, colored according to the percentage of identity. The Viral Quotient (VQ) from pVOG is displayed below or on top of each ORF, with grey meaning that there was no match in the pVOG profiles database and thus no associated VQ value. For prophage #62 (2) that inserted in the core gene *icd*, the boundaries correspond to *icd* on the right and the most distal *att* site found, which is likely to be a remnant prophage border. The most proximal *att* site is also annotated (vertical black line). This figure was generated using the R package GenoPlotR v0.8.9 (Guy et al 2010).

### Klebsiella spp. prophages can be released into the environment

To further characterize the prophages of *Klebsiella pneumoniae* species complex, we sought to experimentally assess their ability to excise and be released into the environment. The analysis described above identified 95 prophages in the 35 strains analyzed from our collection (Figure 3). Among these, 40 were classed as “intact” (including the ten prophages whose genomic composition is characterized in Figure 2). Eleven strains had no “intact” prophages, and four of these also lacked “questionable” prophages. Hence, our collection is representative of the distribution of prophages in the species, with some genomes containing one or multiple putatively functional prophages and others being prophage-free.

**Figure 3.**
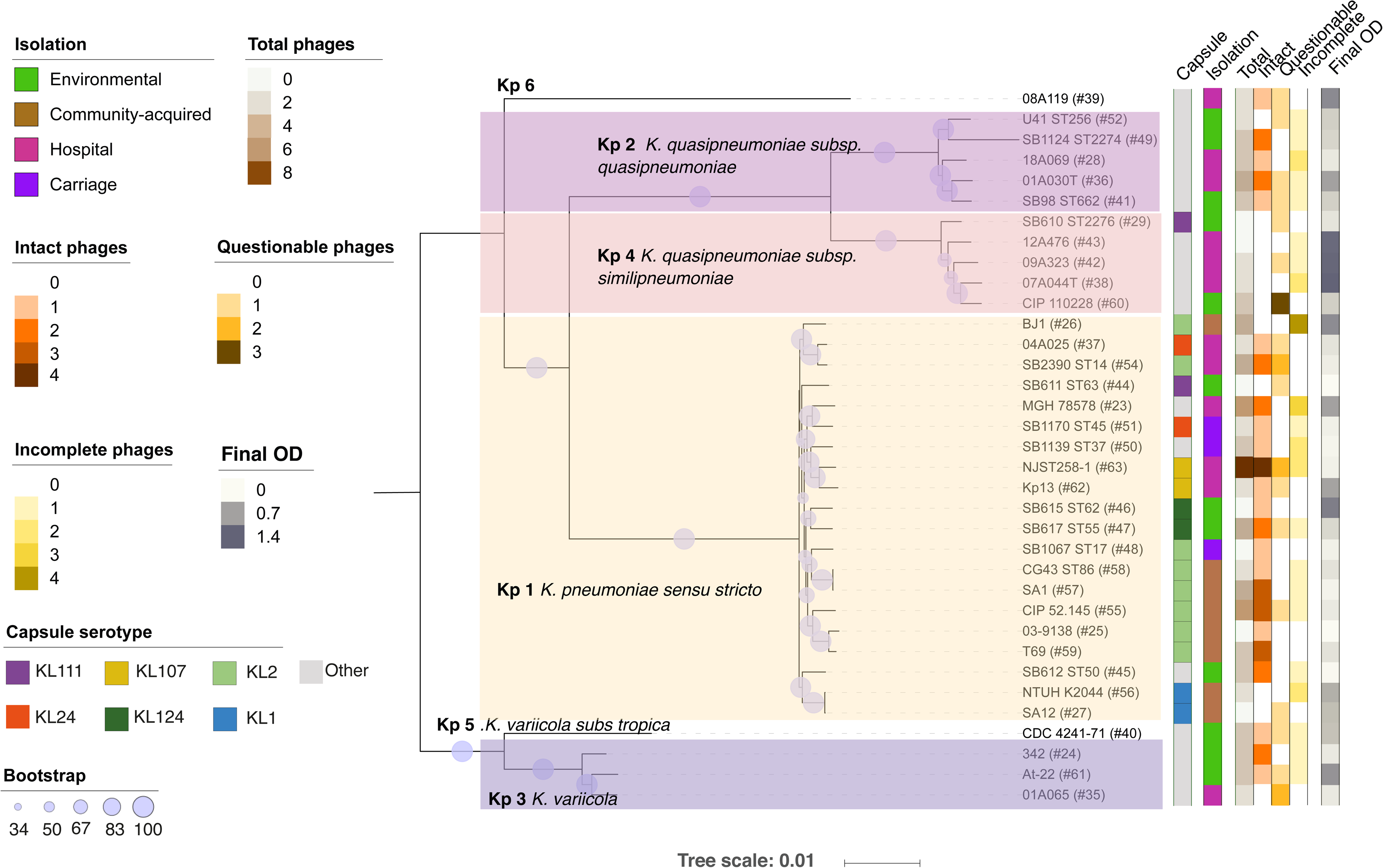
Phylogenetic tree of the 35 *Klebsiella* strains. The tree was built using the protein sequences of 3009 families of the core genome of representative strains from the *Klebsiella pneumoniae* species complex. The first column determines the capsule serotype, the second column provides information about the environment from which it was isolated. The next columns indicate the total, “intact”, “questionable” and “incomplete” prophages detected in the genomes by PHASTER. The last column shows the final absorbance of a culture after induction by mitomycin C. Background colour indicates different *Klebsiella spp*. The size of the circles along the branches are proportional to bootstrap values ranging between 34 and 100. The grey color indicates other CLTs that are only present once in the dataset. # symbol represent the number of the strain in our collection and is used for simplicity throughout the text.

To test prophage induction, we grew these 35 strains and added mitomycin C (MMC) to exponentially-growing cultures. Viable prophages are expected to excise and cause cell death as a result of phage replication and outburst. In agreement with PHASTER predictions, the addition of MMC to the strains lacking “intact” and “questionable” prophages showed no significant cell death (Figure 3 and Figure S4), with the single exception of mild cell death at high doses of MMC for strain NTUH K2044 (#56). Three of the seven strains lacking “intact” but having some “questionable” prophages showed very mild cell death upon induction, two others exhibited a dose-dependent response, and the remaining two showed rapid cell death (#44 and #35). This suggests that at least some of the “questionable” prophages are still inducible and able to kill the host. In addition, all the 24 strains with “intact” prophages showed signs of cell death *ca.* one hour after exposure to MMC (Figure 3 and Figure S4). This occurred in a dose dependent manner, consistent with prophage induction.

These results suggest that most strains have inducible prophages. To verify the release of prophage DNA to the environment, we tested the presence of 17 different phages from eleven different strains by PCR, both in induced and non-induced PEG-precipitated filtered supernatants (Figure S5A). Indeed, all 17 phages were amplified, indicating that viral genomes are released into the environment. As expected, amplification bands were weaker in the non-induced supernatant compared to the induced (Figure S5A). We further verified the recircularization of these prophages in induced supernatants. We detected the recircularization of eight phages induced from six strains (#25, #36, #62, #63, #37 and #50), five of which were classified as “intact”, and three as “questionable” by PHASTER (Figure S5B). Despite using different primers and PCR settings, we did not obtain a clean PCR product for eight phages. This is most likely due to the aforementioned inaccuracies in the delimitation of the prophages, as these phages are detected in induced supernatants (Figure S5A). Overall, our results suggest that most “intact” and some of the “questionable” prophages are able to induce the lytic cycle and lyse the cells.

### The prophage-bacteria interaction network is sparse

Prophages may impact intra-species competition (Bossi et al 2003). In order to characterize this effect in the *Klebsiella pneumoniae* species complex, we produced three independent lysates of all 35 strains, by MMC induction and PEG-precipitation of the filtered supernatants (see Methods). We tested the ability of all lysates to produce a clearing on bacterial overlays of every strain. This resulted in 1225 possible lysate-bacteria combinations, with some lysates being potentially composed of multiple types of phage and different relative proportions across the three independent replicates. We first tested whether the number of different prophages in each strain’s genome correlated with its ability to infect the other strains. To do so, we calculated the infection score of each strain, *i.e.* the average frequency at which its lysate infects the 35 strains of our panel. We observed a positive and significant association between the number of predicted “intact” phages and the infection score (rho=0.16, P < 0.001). Overall, 21 strains out of the 35 (60%) produced a lysate that could infect another strain, but only 75 of the 1225 possible lysate-strain combinations showed clearances or plaques (Figure S6, Figure 4A). To confirm that plaques are caused by the activity of phages, we added lysates to growing cultures of sensitive bacteria, which resulted in bacterial death. Consistent with the action of phages, bacterial death was prevented upon the addition of citrate, a known inhibitor of phage adsorption (Figure S6A). Surprisingly, five out of the 21 strains could be re-infected by their own lysate. This suggests the presence of ineffective mechanisms protecting from super-infection, and is consistent with the observed spontaneous induction of prophages in lysogens in some of these strains (Figure S5, Figure S6C).

**Figure 4.**
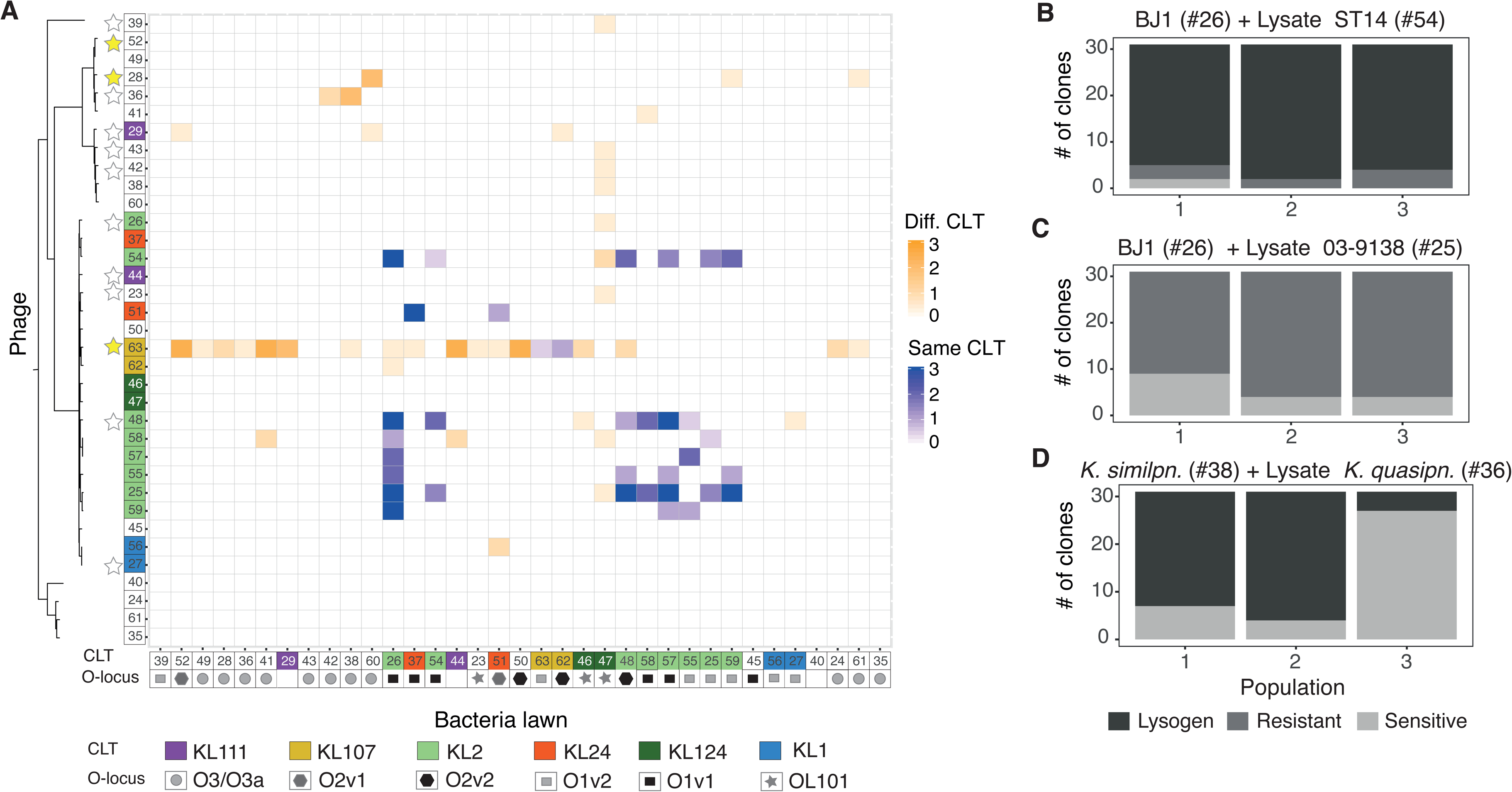
Some *Klebsiella* encode viable prophages that can infect and lysogenize other strains. **A.** Infection matrix indicating the ability of induced and PEG-precipitated supernatants of all strains to form inhibition halos on lawns of all strains. This was repeated three times with three independently-produced lysates. An average of the three experiments is shown. The strains are ordered by phylogeny, and the colours on top of numbers indicate capsule locus types (CLT). Geometric figures along the x-axis indicate the LPS serotype (O-locus). Stars along the y-axis indicate that the genome codes for putative genes involved in colicin production (white) or has a full colicin operon (yellow). Shades of blue indicate infections between lysates produced by bacteria from the same serotype as the target bacteria, whereas shades of orange indicate cross-serotype infections. **B, C and D.** Barplots indicate the number of resistant, sensitive or lysogenized clones from each independent replicate population exposed 24 hours to lysates from other strains. Each bar represents the 31 survivor clones from each of three independent experiments. Grey scale indicates whether: (i) the clones are still sensitive to the phage (light grey), (ii) they became resistant by becoming a lysogen (dark grey) as determined a death upon addition of MMC and PCR targeting prophage sequences, or (iii) they became resistant by other mechanisms.

Some of the inhibition halos observed in the experiments could result from bacteriocins induced by MMC (Ghazaryan et al 2014). To test if this could be the case, we searched for putative bacteriocins in the genomes and found them in 13 of the 35 genomes (Table S1, Figure 4). However, most of them had very low (<50%) protein sequence identity with known proteins and were in genomes lacking recognizable lysis proteins. This may explain why three of the 13 strains failed to produce an inhibition halo on all tested strains and five showed one single mildly inhibited strain (#47). In three of the strains (#63, #48 and #36) we could confirm phage production by recircularization (Figure S5), showing that phages are produced. Indeed, phages from strain #36 can lysogenize strain #38 (see below, Figure 4D), and phages from strain #48 are able to produce plaques on overlays of strain BJ1(Figure S6B). Strain #63 has a full bacteriocin operon, with high identity to colicin E7. It is thus possible that some of the inhibition we observed from the lysates of this strain could be caused by bacteriocins. Note that we also observed that at least two of the prophages of strain #63 are able to excise and circularize (Figure S5 and see annotation Figure 2). Thus, for this particular strain, we cannot precisely identify the cause of the inhibition halos. Finally, from the two remaining strains (#29 and #28, both *K. quasipneumoniae*), only #28 encodes a lysis protein, but they both mildly infect three strains each. Overall, our results suggest that for the majority of the cases shown here, inhibitions seem to be caused by phage activity and are not due to bacteriocins.

Overall, our analysis shows that most lysates cannot infect other strains, resulting in a sparse matrix of phage-mediated competitive interactions. For the 75 phage infections we do observe, only 17 (23%) occurred in all three independently produced lysates (Figure S7), hinting to an underlying stochasticity in the induction and infection process. Interestingly, we observe that infections of target bacteria with lysates from bacteria with the same CLT seem both more reproducible and effective, compared to those lysates produced from bacteria with different CLT (Figure S7), suggesting that the capsule may play a crucial role during phage infection.

Overall, our analysis shows that most lysates cannot infect other strains resulting in a sparse matrix of phage-mediated competitive interactions.

### Induced Klebsiella prophages can lysogenize other strains

An induced prophage can, upon infection of a new host, generate new lysogens. To test the ability of the induced prophages to lysogenize other strains, we challenged *K. pneumoniae* BJ1 (#26) that lacks prophages with lysates from two strains (#54 and #25). These strains have the same CLT as BJ1 (KL2) and we have PCR evidence that they produce viral particles (see above) (Figure S5). We grew BJ1 in contact with these two lysates, and from the surviving BJ1 cells, we isolated 93 clones from 3 independently challenged cultures. We re-exposed these clones to a lysate from strain #54, to test if they had become resistant. Almost all clones (96%) of BJ1 could grow normally upon re-challenge (Figure 4B), suggesting that they had acquired resistance either by lysogenization or some other mechanism. Most of these clones displayed significant cell death upon exposure to MMC (whereas the ancestral strain was insensitive), which is consistent with lysogenization. Analyses by PCR showed that 57 out of 82 clones from the three independent BJ1 cultures exposed to #54 lysate acquired at least one of the “intact” prophages and at least four of them acquired both (shown in Figure 2). In contrast, no BJ1 clones became lysogens when challenged with the lysate of strain #25 (as tested by exposure to MMC and PCR verification, Figure 4C), indicating that the surviving clones became resistant to phage infections by other mechanisms. Overall, this shows that lysogenization of BJ1 is dependent on the infecting phages, with some driving the emergence of alternative mechanisms of resistance to infection in the bacterial host.

To test the taxonomic range of infections caused by induced prophages, we also exposed a Kp 4 (*K. quasipneumoniae subsp similipneumoniae*) (#38) to a lysate from a Kp 2 (*K. quasipneumoniae subsp quasipneumoniae*) (#36) that consistently infects #38 even if it belongs to a different subspecies and has a different capsular CLT (Figure 4A). Survivor clones of strain #38 were lysogenized (in 61 out of 93 surviving clones) exclusively by one of the phages present in strain #36, as confirmed by PCR of the phage genome (Figure 4D). Interestingly, four of these lysogenized clones did not display cell death when exposed to MMC, suggesting that their prophages might not be fully functional. Taken together, our results show that some *Klebsiella* prophages can be transferred to and lysogenize other strains, including from other subspecies.

### Resistance to super-infection and bacterial defences do not explain the interaction matrix

To investigate the determinants of infections in the interaction matrix, we first hypothesized that resident prophages could hamper infections by lysates of other strains. Consistently, the most sensitive strain in our panel (#26) does not have any detectable prophages, rendering it sensitive to infection by numerous lysates. However, we found no negative association between the number of prophages in the target strain and the number of times it was infected (rho=0.04, P= 0.166). Repression of infecting phages is expected to be highest when there are similar resident prophages in the strain. Even if closer strains are more likely to carry the same prophages, the interaction matrix clearly shows that phages tend to infect the most closely related strains. To determine if the presence of similar prophages shapes the infection matrix, we calculated the genetic similarity between all “intact” prophages using weighted Gene Repertoire Relatedness (wGRR) (see Methods). The frequency of infection was found to be independent from the similarity between prophages (determined as higher than 50% wGRR for phage pairs, Odds ratio = 0.55, P-value= 0.055) when analysing only “intact” prophages or also including the “questionable” (Figure S8A). Ultimately, resident phages may repress incoming phages if they have very similar repressors, but pairs of dissimilar phages (wGRR<50%) with similar repressors (>80% sequence identity) are only ∼ 0.3% of the total (see Methods, Figure S8B). Thus, the analysis of both phage sequence similarity and similarity between their repressor suggests that the observed interaction matrix is not strongly influenced by the resistance to super-infection provided by resident prophages.

Bacterial defence systems block phage infections and are thus expected to influence the interaction matrix. To study their effect, we analysed systems of CRISPR-Cas and restriction-modification (R-M), since these are the most common, the best characterized to date, and those for which tools for their computational detection are available (Labrie et al 2010, Samson et al 2013). We first searched for these systems in the bacterial genomes. We found CRISPR-Cas systems in 8 of the 35 strains and tested if the strains that were infected by the lysates of other bacteria were less likely to encode these systems. We observed no correlation between the number of unsuccessful infections and the presence of CRISPR-Cas systems in a genome (P> 0.05, Wilcoxon test). Accordingly, the majority of strains (77%) lack spacers against any of the prophages of all the other strains and we found no correlation between the presence of CRISPR spacers targeting incoming phages and the outcome of the interaction (i.e., infection or not) (Odds-ratio= 1.28 Fisher’s exact test, P-value = 1, Dataset S1, Figure S9A). Actually, only 5% of the pairs with null interactions (no infection) concerned a target strain with CRISPR spacers matching a prophage in the lysate-producing strain. This indicates that CRISPR-Cas systems do not drive the patterns observed in the infection matrix.

The analysis of R-M systems is more complex because their specificity is often harder to predict. We searched for these systems in the 35 genomes and identified the following systems: 37 type I, 158 type II, 4 type IIG, 7 type III and 7 type IV. We then investigated whether the systems present in the target strains could protect from phages in the lysates. Almost all the strains studied here have systems that could target at least one prophage from the lysate-producing strains (Figure S9B, see Methods). However, the joint distribution of R-M systems and targeted prophages does not explain the outcome observed in the infection matrix (Odds-ratio= 0.78, Fisher’s exact test on a contingency table, P-value = 0.4, Figure S9). We thus conclude that the distribution of either CRISPR-Cas or R-M bacterial defence systems in the bacterial strains targeted by the phages in the lysates does not explain the network of phage-bacteria interactions we observe.

### The capsule plays a major role in shaping phage infections

The first bacterial structures that interact with phages are the capsule and the LPS. We thus tested whether their serotypes shape the infection network of the *Klebsiella* prophages. We used Kaptive to serotype the strains from the genome sequence and tested if the infections were more frequent when the target bacteria and the one producing the lysate were from the same serotype. We found no significant effect for the LPS serotypes (Figure 4A, Fisher’s exact test P=0.37). In contrast, we observed 35 cross-strain infections between lysates and sensitive bacteria from the same CLT out of 105 possible combinations (33%), whereas only 3.6% of the possible 1120 inter-CLT infections were observed (40 infections in total, Figure 4A, P<0.0001, Fisher’s exact test). For example, strain #37 was only infected by lysates produced by the other strain with the KL24 CLT (#51) (Figure 2). Similarly, strains #57, #55 and #25 were only infected by lysates from strains of the same CLT (KL2) (Figure 4A). These results are independent of the genetic relatedness, as we observed infections between strains with the same CLT that are phylogenetically distant. Intriguingly, the lysate of one strain alone (#63) produced an inhibition halo in lawns of fifteen strains from different CLTs, but (as mentioned above) this may be caused by a bacteriocin present in this strain’s genome. To control for the putative effect of bacteriocins in the association between infections and the CLT, we restricted the analysis to strains lacking bacteriocin homologs (N = 22) and those with a CLT determined with high confidence by Kaptive (N = 29) (see Methods). We confirmed in both cases that CLT is positively associated with cross-strain infections (P<0.0001, Fisher’s exact test for both).

The specificity of induced prophages for strains with a CLT similar to the original cell could be explained by the presence of depolymerases in their tail fibres. Visual observation of plaques showed that the latter are small and are not surrounded by enlarged halos typically indicative of depolymerase activity (Figure S6). We also searched the genomes of the prophages for published depolymerase domains using protein profiles and a database of known depolymerases (see Methods, Table S2). We only found hits to two protein profiles (Pectate_lyase_3 and Peptidase_S74) with poor e-value (> 10^-10^) and a low identity homolog (< 50%) to one protein sequence (YP_009226010.1) (Table S3 and S4). Moreover, these homologs of depolymerases were found in multiple prophages that are hosted in strains differing in CLTs, suggesting that they are either not functional, or that they do not target a specific CLT. It is also possible that the CLT has switched (*ie* changed by recombination (Wyres et al 2015)) since the phage originally infected the strain. This suggests that the depolymerases present in these prophages (or at least those that we could identify) do not explain the patterns observed in the interaction matrix, raising the possibility that novel depolymerases remain to be found.

It is often mentioned in the literature that capsules protect against phages (Scholl et al 2005). However, the results above show that induced prophages tend to infect strains of similar CLT, and capsule mutants of the strain Kp36 were previously shown to become resistant to the virulent phage 117 (Tan et al 2020). Hence, we wondered if strains lacking a capsule would also be immune to temperate phages,, and so we tested whether isogenic capsule mutants from different CLTs (KL2, KL30 and KL1) were sensitive to phages present in the lysates. We verified by microscopy and biochemical capsule quantification that these mutants lacked a capsule and then challenged them with the lysates of all 35 strains. Wild type strains #56 (K1) and #24 (K30) are resistant to all lysates, as are their capsule mutants. Capsule mutants from #26 (BJ1), #57 and #58 (all KL2), and #37 (KL24) were resistant to phages, whilst their respective wild-types are all sensitive to phage infections (Figure 5). Hence, phages targeting these strains only infect when a capsule is present. These results show that the loss of the capsule does not make bacteria more sensitive to phages. On the contrary, the capsule seems required for infection.

**Figure 5.**
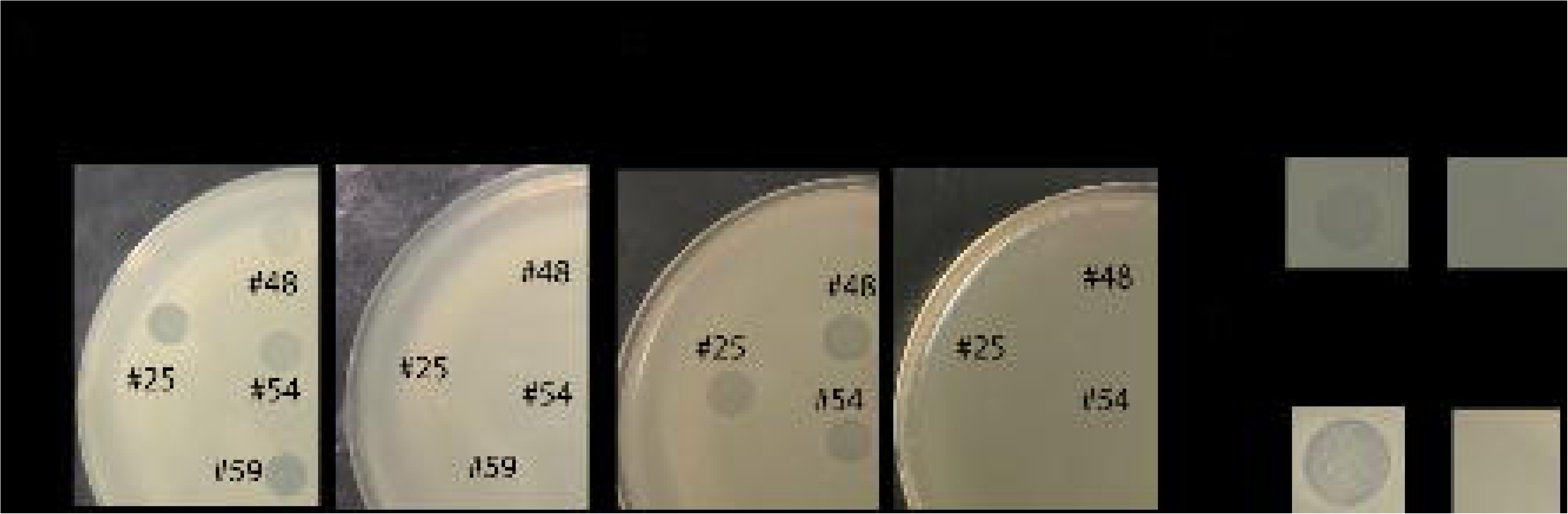
The capsule is required for phage infection. Lawns of different strains, wild type (Cap +) and capsule null mutant (Cap-) in contact with PEG-precipitated supernatant of other strains. Isogenic capsule mutants of strains #26 and #37 were constructed by an in-frame deletion of *wza* gene, and a *wcaJ* deletion for strains #57 and #58 (see Methods).

### Phage predation stimulates capsule loss

Given that non-capsulated mutants are resistant to phages (as shown above and also observed in other studies (Tan et al 2020)), we hypothesized that phages that infect a particular strain could drive the loss of its capsule. To test this, we performed a short-term evolution experiment (ten days, ca 70 generations). We assessed the emergence of non-capsulated bacteria in 3 different strains: (i) a strain with no temperate phages (BJ1: #26); (ii) a strain that produces phages infecting many strains including itself (#54) even in the absence of MMC induction (Figure S6C); and (iii) a strain that is resistant to its own phages (#36). Both strains #36 and #54 produce infectious phages which can lysogenize other strains (#38 and #26, respectively, Figure 4B). Six independent populations of each strain were evolved in four different environments: (i) LB, (ii) LB supplemented with 0.2% citrate to inhibit phage adsorption, (iii) LB with MMC to increase the phage titre, and (iv) LB with MMC and 0.2% citrate to control for the effect of faster population turnover due to prophage induction and the subsequent cellular death. We expected that passages in rich medium might lead to non-capsulated mutants, even in the absence of phages, as previously observed (Buffet et al 2020, Julianelle 1928, Randall 1939). This process should be accelerated if phage predation drives selection of the non-capsulated mutants (presence of phages, absence of citrate, in LB). It should be further accelerated if the intensity of phage predation increases (under MMC). As expected, all strains progressively lost their capsule albeit at different rates (Figure 6A). To allow comparisons between treatments and strains, we calculated the area under the curve during the first five days, where most of the capsule loss took place (Figure 6B). The strain BJ1 lacks prophages and shows no significant differences between the treatments. Strain #54 lost its capsule faster when prophages were induced (MMC) and citrate relieved this effect in accordance with the hypothesis that the speed of capsule loss depends on the efficiency of phage infection (MMC+citrate). Similarly, in the treatment with citrate the capsule is lost at a slower rate than in the LB treatment. In the latter, the few events of spontaneous prophage induction generate a basal level of predation that is sufficient to increase the rate of loss of the capsule (albeit not to the levels of the MMC treatment). Note that the combination of citrate and MMC did not significantly affect bacterial growth, and thus the effect we observe seems to be caused by phage predation (Figure S6 and Figure S10). Finally, strain #36 showed no significant difference between the experiments with MMC and LB. This suggests that the amount of phages in the environment does not affect the rate of capsule loss in this strain, consistent with it being insensitive to its own phage. Intriguingly, adding citrate lowered the rate of capsule loss in this strain, a result that suggests that even if phage infection is inefficient, there may be small deleterious impact on the presence of phages. Taken together, these results show that effective predation by induced prophages selects for the loss of the capsule in the lysogen.

**Figure 6.**
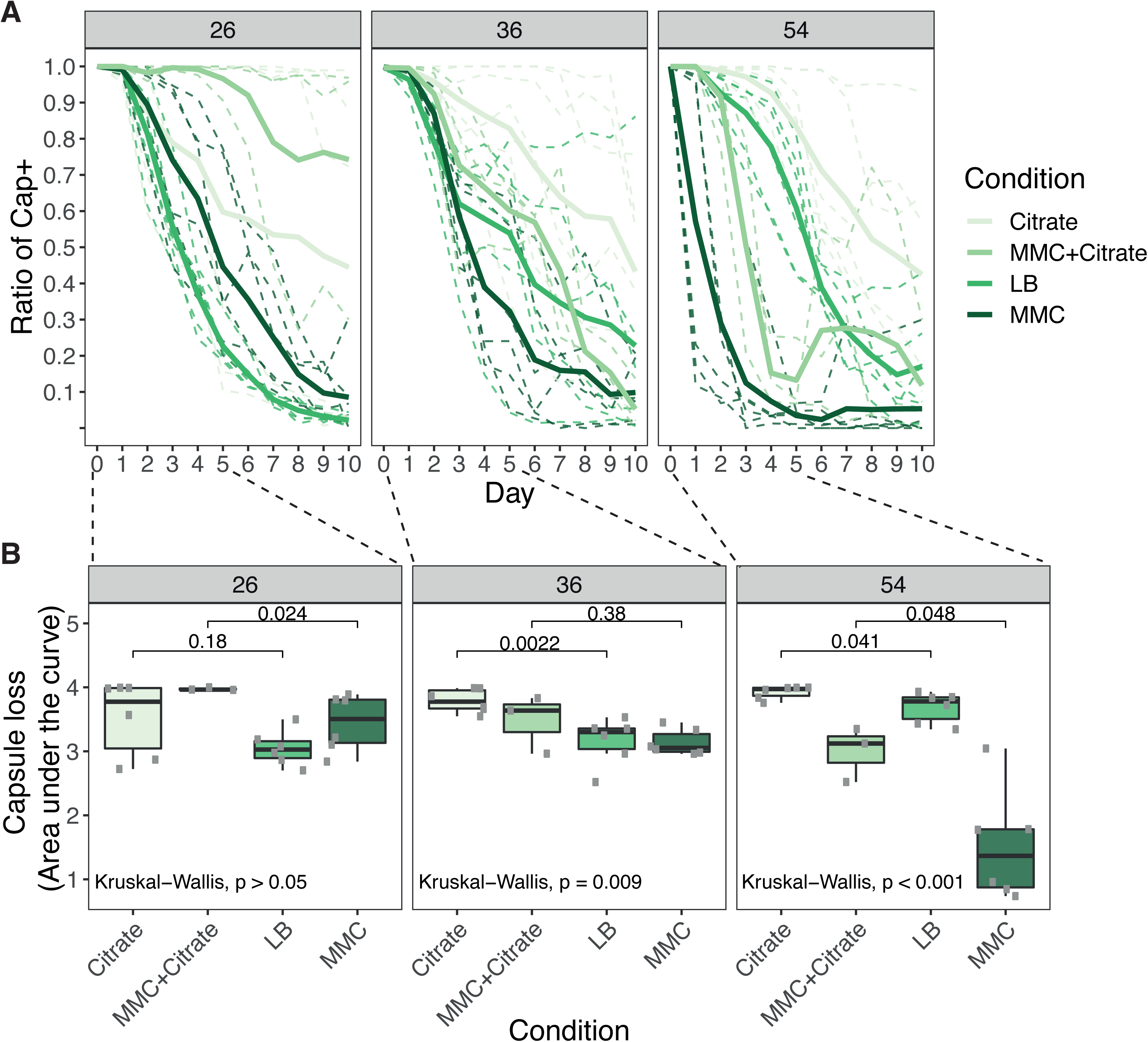
Loss of capsule in three *Klebsiella* strains. **A.** Ratio of capsulated clones throughout the ten days before daily passages of each culture. Shades of green represent the different environments in which evolution took place. MMC stands for mitomycin C. Full lines represent the average of the independent populations of the same strain and environment (shades of green). Dashed lines represent each of the independent populations. **B**. Area under the curve during the first five days of the experiment.

## DISCUSSION

We provide here a first comprehensive analysis of the distribution of prophages in *Klebsiella* genus, their genetic composition and their potential to excise, infect and lysogenize other strains. The number of prophages varies significantly across the species of the genus, but most genomes of *Klebsiella* encode for a phage or its remnants. *K. pneumoniae* is one of the species with more prophages among widely sequenced bacteria, suggesting that temperate phages are particularly important for its biology. This rapid turnover of prophages, already observed in other species, may contribute to the phenotypic differences between strains. Since many of these prophages seem to retain the ability to excise, form viable virions, and lysogenize other bacteria, they could spur adaptation when transferring adaptive traits (including antibiotic resistance in clinical settings). Further work will be needed to assess the phenotypic consequences of lysogeny and the frequency of transduction, but these results already show that the contribution of prophages to *Klebsiella* genomes is significant.

Phage-bacteria interactions shape a myriad of biological processes. Several recent studies have detailed infection networks to understand the ecological traits and molecular mechanisms shaping them (Flores et al 2011, Mathieu et al 2020, Weitz et al 2013). These analyses use isolated phages from laboratory stocks, or phages recovered from co-evolution experiments (Flores et al 2011). Additionally, these studies tend to focus on virulent phages rather than temperate, either because they envision some sort of phage therapy or because virulent phages give simpler phenotypes. Several recent computational studies have described the different phage families and the beneficial traits they may impart to their hosts, such as virulence factors (Bobay et al 2013, Brueggemann et al 2017, Castillo et al 2018, Mathieu et al 2020). A few have explored the natural diversity of temperate phages of a species and experimentally assessed their ability to cross-infect other strains (Mathieu et al 2020). Here, we sought to generate a network of temperate phage-mediated bacterial interactions in naturally infected *Klebsiella* spp, including competitive killing through phage induction and also the transfer of prophages between strains. These results are especially pertinent for *K. pneumoniae* in clinical settings because many antibiotics stimulate prophage induction (Allen et al 2011, Otsuji et al 1959, Wagner and Waldor 2002) and facilitate phage infection (Comeau et al 2007, Kim et al 2018). Also, phages in the mammalian gut, the most frequent habitat of *K. pneumoniae*, tend to be temperate and result from *in situ* prophage induction (De Paepe et al 2016, Mathieu et al 2020).

We studied a large cohort of strains from the *K. pneumoniae* species complex, representing a considerable diversity in terms of the number and types of prophages present in lysogenic strains. However, the extrapolation of some of the results presented here to the *Klebsiella* genus as a whole must be done with care. Further work, and a larger picture, will be possible once genomes from other clades of the genus are available. Moreover, since it would be unfeasible to generate mutants for all the strains, or perform evolution experiments with the entire cohort, some experimental assays performed here were limited to a reduced number of strains and environments. It is thus possible that some observations may not be consistently obtained with other genetic backgrounds, or ecological contexts (e.g., spatially structured environments). Nevertheless, our work considers a total of 1225 lysate-bacteria interactions, which already allows the characterization of a wide range of possible outcomes.

Our experimental setup was specifically designed to address the natural variation in the ability of lysates to infect other strains. In order to capture this variation, the network of infections was performed thrice with three independently generated lysates, which led to stochastic variation in the infection outcomes. This has been rarely characterized before for poly-lysogens. One possible explanation for the observed stochasticity in the outcomes of infections could be that the resident phages are unfit due to the accumulation of deleterious mutations during lysogeny. This could affect, for instance, the number of produced virions, and thus the infection efficiency. Alternatively, there could be a noteworthy degree of natural variation in the frequencies of each induced phage across the independently-generated lysates. This means that the underrepresentation of certain phages could lead to unsuccessful infections. Having multiple prophages is expected to be costly (in terms of gene expression and cell death by induction), but extends the range of competitors that can be affected by prophage induction. Finally, infections of target bacteria with lysates from bacteria with the same CLT seem both more reproducible and effective, compared to those lysates produced from bacteria with different CLT (Figure S7), suggesting that stochasticity in infections can also be a consequence of capsule-phage interactions, e.g. if the former is unevenly distributed throughout the cell envelope (Phanphak et al 2019) or if there is stochasticity in its expression (Krinos et al 2001, Tzeng et al 2016). Future experiments tackling diverse ecological scenarios (e.g., co-cultures between three or more strains) will help understanding the causes and consequences of multiple infections producing poly-lysogens for competitive and evolutionary interactions between strains.

Overall, we found that 60% of the strains could produce at least one lysate that led to phage infection of at least one other strain. The number of strains able to produce phages is comparable to what is observed in other species like *Pseudomonas aeruginosa* (66%) (Bondy-Denomy et al 2016) and *Salmonella enterica* (68%) (Zhang and LeJeune 2008). However, we only observed 6% of all the possible cross-infections. This is much less than in *P. aeruginosa,* where a set of different lysogenic strains derived from PA14 was infected by *ca* 50% of the lysates tested (Bondy-Denomy et al 2016). Similarly, in *S. enterica,* ∼35% of cross-infections were effective (Zhang and LeJeune 2008). This suggests that the likelihood that a prophage from one strain is able to infect another *Klebsiella* strain is relatively small. Hence, when two different *Klebsiella* strains meet, they will be very often immune to the prophages of the other strains. This implicates that prophages would be less efficient in increasing the competitive ability of a *Klebsiella* strain than in other species. Finally, this also implies that phage-mediated HGT in *Klebsiella* may not be very efficient in spreading traits across the species, which means that, in some situations, the capsule could slow down evolution by phage-mediated horizontal gene transfer.

Interestingly, 5 out of 35 strains can be infected by their own induced prophages. It is commonly assumed that lysogens are always resistant to re-infection by the same phage (Brussow and Kutter 2005), but the opposite is not unheard of. It has been previously reported that *E. coli* strains could be re-infected by the same phage twice, both by phage lambda (Calef et al 1965) and by phage P1 (Scott et al 1978). At this time, we can only offer some speculations for this lack of superinfection immunity. For example, it has been estimated that there are only 10 free dimmers of Lambda’s cI repressor in a typical cell (Bakk and Metzler 2004). The infection of several phages at the same time may titre this limited amount of repressor, such that some incoming phages are able to induce resident prophages to enter a lytic cycle. This could be amplified when prophages accumulate deleterious mutations in the repressor or in the binding sites of the repressor, further decreasing its ability to silence the prophage or the incoming phage. This could also explain why we observe frequent prophage spontaneous induction (strain #54, Figure S6C).

The small number of cross-infectivity events could not be attributed to bacterial defence systems (R-M or CRISPR-Cas) or LPS serotypes. The similarity between prophages or their repressor proteins, potentially facilitating superinfection immunity (Bondy-Denomy et al 2016), also failed to explain the observed infection patterns. Instead, the relatively few cases of phage infection in the matrices are grouped in modules that seem determined by the capsule composition, since there is a vast over-representation of infections between bacteria of the same CLT. This seems independent of their genetic relatedness, since we observed infections between strains with the same CLT that are phylogenetically distant. This CLT specificity is consistent with a recent study that focuses on a carbapenemase-producing *K. pneumoniae* CG258, in which very few phages could lyse bacteria with different CLT (Venturini et al 2019). The CLT-specificity of phage-encoded depolymerases (Hsieh et al 2017, Lin et al 2014, Majkowska-Skrobek et al 2018, Niemann et al 1977a, Niemann et al 1977b, Pan et al 2017, Pan et al 2019, Thurow et al 1974) could explain these results. Yet, in our set, few of such enzymes were detected, they exhibited very low identity to known depolymerases, and did not seem to correlate with the CLT. At this stage, it is difficult to know if depolymerases are rare in temperate phages or if they are just too different from known depolymerases, since these were mostly identified from virulent phages. If depolymerases are indeed rare in temperate phages, this raises the important question of the alternative mechanisms that underlie their capsule specificity.

Most of the literature on other species concurs that the capsule is a barrier to phages by limiting their adsorption and access to cell receptors (Moller et al 2019, Negus et al 2013, Scholl et al 2005). In contrast, our results show that the capsule is often required for infection by *Klebsiella* induced prophages. This is in agreement with a recent study where inactivation of *wcaJ*, a gene essential for capsule synthesis, rendered the strain resistance to phage infection by a virulent phage (Tan et al 2020). Specific interactions between the phage and the capsule could be caused by the latter stabilizing viral adsorption, allowing more time for efficient DNA injection (Bertozzi Silva et al 2016). This may have resulted in phages selecting for the ability to recognize a given capsule serotype. The specificity of the interactions between temperate phages and bacteria, caused by capsule composition, has outstanding implications for the ecology of these phages because it severely limits their host range. Indeed, we report very few cases (*ca*. 3%) of phages infecting strains with different CLTs, and this could be an overestimation if we discard strain #63 because of its potential bacteriocin activity. The consequence of this evolutionary process is that phage pressure results in selection for the loss of capsule because non-capsulated bacteria are resistant to phage infection. Our experimental evolution captures the first steps of this co-evolution dynamic, suggesting that phage predation selects very strongly for capsule loss in the infected strains. Since capsules are prevalent in *Klebsiella,* this suggests that it may be re-acquired later on.

Our results might also provide insights regarding the possible use of phages to fight the increasing challenge of antibiotic resistance in *Klebsiella* infections. Although more work is needed to understand how to best use virulent phages to control *Klebsiella* infections, our results already hint that phage therapy may, at least in a first step, lead to capsule loss. While such treatments may be ineffective in fully clearing *Klebsiella* infections (due to the large diversity of existing serotypes), they can select for non-capsulated mutants. The latter are expected to be less virulent (Paczosa and Mecsas 2016), because the capsule is a major virulence factor in *K. pneumoniae*. The increase in frequency of non-capsulated mutants may also increase the efficiency of traditional antimicrobial therapies, as the capsule is known to increase tolerance to chemical aggressions, including antibiotics and cationic antimicrobial peptides produced by the host (Paczosa and Mecsas 2016).

## MATERIALS AND METHODS

### Strains and growth conditions

We used 35 *Klebsiella* strains that were selected based on based on MLST data, and representative of the phylogenetic and clonal diversity of the *Klebsiella pneumoniae species* complex (Blin et al 2017). Strains were grown in LB at 37° and under shaking conditions (250 rpm).

### Genomes

254 genomes of *Klebsiella* species (of which 197 of *K. pneumoniae*) and one *Cedecea* sp (outgroup) were analysed in this study. This included all complete genomes belonging to *Klebsiella* species from NCBI, downloaded February 2018, and 29 of our own collection(Blin et al 2017). We corrected erroneous NCBI species annotations using Kleborate typing (Wyres et al 2016). 253 genomes were correctly assigned to its species, with a “strong” level of confidence, as annotated by Kaptive. Only two genomes were classified with a “weak” level of confidence by the software. All information about these genomes is presented in Dataset S1.

### Identification of prophages

To identify prophages, we used a freely available computational tool, PHASTER (Arndt et al 2016), and analysed the genomes on September 2018. The completeness or potential viability of identified prophages are identified by PHASTER as “intact”, “questionable” or “incomplete” prophages. All results presented here were performed on the “intact” prophages, unless stated otherwise. Results for all prophages (“questionable”, “incomplete”) are presented in the supplemental material. Primers used for phage detection and recircularization are presented in Table S5.

### Prophage characterization

*i. Prophage delimitation*. Prophages are delimited by the *attL* and *attR* recombination sites used for phage re-circularization and theta replication. In some instances, these sites were predicted by PHASTER. When none were found, we manually searched for them either by looking for similar *att* sites in related prophages or by searching for interrupted core bacterial genes. ii. *Functional annotation* was performed by combining multiple tools: *prokka* v1.14.0 (Seemann 2014), pVOG profiles (Grazziotin et al 2017) searched for using *HMMER* v3.2.1 (Eddy 2011), the PHASTER Prophage/Virus DB (version Aug 14, 2019), *BLAST 2.9.0+* (Camacho et al 2009), and the *ISFinder* webtool (Siguier et al 2006). For each protein and annotation tool, all significant matches (e-value<10^-5^) were kept and categorized in dictionaries. If a protein was annotated as “tail” in the description of a matching pVOG profile or PHASTER DB protein, the gene was categorized as tail. Results were manually curated for discrepancies and ties. For proteins matching more than one pVOG profile, we attributed the Viral Quotient (VQ) associated to the best hit (lowest e-value). The Viral Quotient (VQ) is a measure of how frequent a gene family is present in phages, and ranges from zero to one with higher values meaning that the family is mostly found in viruses. *iii. Repressor identification*. Repressors in the prophages were detected using specific HMM protein profiles for the repressor, available in the pVOG database (Grazziotin et al 2017). We selected those with a VQ higher than 0.8. These profiles were matched to the *Klebsiella* intact prophage sequences using HMMER v3.1b2, and we discarded the resulting matches whose best domain had an e-value of more than 0.0001 or a coverage of less than 60%. We further selected those with at least one of the following terms in the descriptions of all the proteins that compose the HMM: immunity, superinfection, repressor, exclusion. This resulted in a set of 28 profiles (Table S6). The similarity between repressors was inferred from their alignments (assessed with the *align.globalxx* function in from the *pairwise2* module in biopython, v1.74), by dividing the number of positions matched by the size of the smallest sequence (for each pair of sequences).

### Core genome

i. *Klebsiella spp*. (*N*=255) (Figure 1A and S1). The core genome was inferred as described in (Touchon et al 2014b). Briefly, we identified a preliminary list of orthologs between pairs of genomes as the list of reciprocal best hits using end-gap free global alignment, between the proteome of a pivot and each of the other strains proteome. Hits with less than 80% similarity in amino acid sequences or more than 20% difference in protein length were discarded. ii. *Laboratory collection of Klebsiella genomes* (*N*=35) (Figure 4). The pangenome was built by clustering all protein sequences with Mmseqs2 (v1-c7a89) (Steinegger and Soding 2017) with the following arguments: *-cluster-mode 1* (connected components algorithm), *-c 0.8 –cov-mode 0* (default, coverage of query and target >80%) and *–min-seq-id 0.8* (minimum identity between sequences of 80%). The core genome was taken from the pan-genome by retrieving families that were present in all genomes in single copy.

### Phylogenetic trees

To compute both phylogenetic trees, we used a concatenate of the multiple alignments of the core genes aligned with MAFFT v7.305b (using default settings). The tree was computed with IQ-Tree v1.4.2 under the GTR model and a gamma correction (GAMMA). We performed 1000 ultrafast bootstrap experiments (options *–bb* 1000 and *– wbtl*) on the concatenated alignments to assess the robustness of the tree topology. The vast majority of nodes were supported with bootstrap values higher than 90% (Figure S1). The *Klebsiella spp.* (N=255) tree had 1106022 sites, of which 263225 parsimony-informative. The phylogenetic tree of the laboratory collection strains (N = 35) was built using 2800176 sites, of which 286156 were parsimony-informative.

### Capsule serotyping

We used Kaptive, integrated in Kleborate, with default options (Wyres et al 2016). Serotypes were assigned with overall high confidence levels by the software (see Dataset S1): 13 were a perfect match to the sequence of reference, 144 had very high confidence, 33 high, 35 good, 11 low and 19 none. From the strains used in the experiments, two were a perfect match to reference strains, 20 had very high confidence, 3 were high, 7 good and 3 for which the assignment had low confidence.

### Bacteriocin detection

We checked for the presence of bacteriocins and other bacterial toxins using BAGEL4 (van Heel et al 2018). The results are reported in Table S1.

### Depolymerase detection

We checked for the presence of 14 different HMM profiles associated with bacteriophage-encoded depolymerases from multiple bacterial species (Pires et al 2016), Table S2). The profiles were matched against the complete set of prophage proteins using HMMERv3.1b2, filtering by the e-value of the best domain (maximum 10^-3^) and the coverage of the profile (minimum 30%). Five additional sequences of depolymerases, validated experimentally in lytic phages of *Klebsiella* (see references in Table S1), were also matched against our dataset using *blastp* (version 2.7.1+, default parameters, and filtering by the e-value (maximum 10^-5^), identity (40%) and coverage (40%).

### Identification of CRISPR arrays and R-M systems

(i) *CRISPR-Cas arrays*. We used CRISPRCasFinder (Couvin et al 2018) (v4.2.18, default parameters) to identify the CRISPR arrays in all *Klebsiella* genomes used in this study. We excluded arrays with less than 3 spacers. We then matched each spacer sequence in each array with the complete prophage nucleotide sequences *(blastn* version 2.7.1+, with the -task blastn-short option). Only matches with a spacer coverage of at least 90%, a maximum e-value of 10^-5^ and a minimum nucleotide identity of 90% were retained. The resulting matches indicate prophages that are targeted by these spacers, and the full set of these results are presented in Dataset S1. (ii) *R-M systems* were identified using the highly specific and publicly available HMM profiles in https://gitlab.pasteur.fr/erocha/RMS_scripts. If a single protein matched multiple systems, the best hit for each protein-R-M pair was selected. To assess the likelihood that a strain can defend itself against infection by prophages induced from another strain using R-M systems, we calculated the similarity between these proteins (all versus all) using BLAST (*blastp* version 2.7.1+, filtering by identity > 50%, e-value < 0.0001). We then considered that two R-M associated proteins can target similar recognition sites if their identity is either at above 50% (for Type II and IV REases), 55% (for Type IIG), 60% (for Type II MTases)) or 80% (for Type I and III MTases and Type I and III REases), according to (Oliveira et al 2016). We inferred the recognition sites targeted by these systems using REBASE (http://rebase.neb.com/rebase/rebase.html, default parameters). We selected the recognition site associated to the protein with the best score, and further chose those whose identity obeyed the thresholds enumerated above (for each type of RMS). The nucleotide motifs associated with these recognition sites were searched for in the intact prophage genomes from the 35 bacterial strains using the Fuzznuc program from the EMBOSS suite (version 6.6.0), with the default parameters, using the option to search the complementary strand when the motif was not a palindrome. Finally, for each pair of bacterial strains A-B (where strain A produces the lysate and strain B is used as a bacterial lawn), we looked for prophages in strain A whose genomes contain recognition sites targeted by R-M systems present in strain B and absent from strain A. If a recognition site was found in any prophage from strain A, we consider that prophage to be putatively targeted by (at least) one R-M defence system of strain B. Because some R-M systems require two (similar) recognition sites in order to effectively target incoming DNA (Mucke et al 2003), we also did a separate analysis where we require that each motif is found twice in each individual prophage genome. This analysis resulted in similar qualitative results.

### Prophage experiments

*(i) Growth curves:* 200 µL of diluted overnight cultures of *Klebsiella spp*. (1:100 in fresh LB) were distributed in a 96-well plate. Cultures were allowed to reach OD_600_ = 0.2 and either mitomycin C to 0, 1 or 3 µg/mL or PEG-precipitated induced and filtered supernatants at different PFU/ml was added. Growth was then monitored until late stationary phase. *(ii) PEG-precipitation of virions*. Overnight cultures were diluted 1:500 in fresh LB and allowed to grow until OD_600_ = 0.2. Mitomycin C was added to final 5 µg/mL. In parallel, non-induced cultures were grown. After 4h hours at 37°C, cultures were centrifuged at 4000 rpm and the supernatant was filtered through 0.22um. Filtered supernatants were mixed with chilled PEG-NaCl 5X (PEG 8000 20% and 2.5M of NaCl) and mixed through inversion. Virions were allowed to precipitate for 15 min and pelleted by centrifugation 10 min at 13000 rpm at 4°C. The pellets were dissolved in TBS (Tris Buffer Saline, 50 mM Tris-HCl, pH 7.5, 150 mM NaCl). *(iii) All-against-all infection*. Overnight cultures of all strains were diluted 1:100 and allowed to grow until OD_600_ = 0.8. 1mL of bacterial cultures were mixed with 12 mL of top agar (0.7% agar), and 3 mL of the mixture was poured onto a prewarmed LB plate and allowed to dry. 10 µL of freshly prepared and PEG-precipitated lysates were spotted on the top agar and allowed to grow for 4h at 37° prior to assessing clearance of bacterial cultures. This was repeated in three independent temporal blocks. *(iv) Calculating plaque forming units (PFU)*. Overnight cultures of sensitive strains were diluted 1:100 and allowed to grow until OD_600_ = 0.8. 250 µL of bacterial cultures were mixed with 3 mL of top agar (0.7% agar) and poured intro prewarmed LB plates. Plates were allowed to dry before spotting serial dilutions of induced and non-induced PEG-precipitate virions. Plates were left overnight at room temperature and phage plaques were counted.

### Evolution experiment

Three independent clones of each strain (#54, #26 and #36) were used to initiate each evolving of the three evolving populations in four different environments: (i) LB, (ii) LB supplemented with 0.2% citrate (iii) LB with mytomycin C (MMC, 0.1 µg/mL) and (iv) LB with MMC (0.1 µg/mL) and supplemented with 0.2% citrate. Populations were allowed to grow for 24h at 37°. Each day, populations were diluted 1:100 and plated on LB to count for capsulated and non-capsulated phenotypes. This experiment was performed in two different temporal blocks and its results combined.

### wGRR calculations and network building

We searched for sequence similarity between all proteins of all phages using mmseqs2 (Steinegger and Soding 2017) with the sensitivity parameter set at 7.5. The results were converted to the blast format for analysis and were filtered with the following parameters: e-value lower than 0.0001, at least 35% identity between amino acids, and a coverage of at least 50% of the proteins. The filtered hits were used to compute the set of bi-directional best hits (bbh) between each phage pair. This was then used to compute a score of gene repertoire relatedness for each pair of phage genomes, weighted by sequence identity, computed as following:

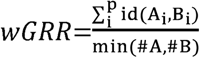

where A_i_ and B_i_ is the pair *i* of homologous proteins present in *A* and *B* (containing respectively #A and #B proteins and *p* homologs), id(*A_i_*,*B_i_*) is the percent sequence identity of their alignment, and min(#*A*,#*B*) is the total number of proteins of the smallest prophage, *i.e.* the one encoding the smallest number of proteins (*A* or *B*). wGRR varies between zero and one. It amounts to zero if there are no orthologs between the elements, and one if all genes of the smaller phage have an ortholog 100% identical in the other phage. Hence, the wGRR accounts for both frequency of homology and degree of similarity among homologs. For instance, three homologous genes with 100% identity between two phages, where the one with the smallest genome is 100 proteins long, would result in a wGRR of 0.03. The same wGRR value would be obtained with six homologous genes with 50% identity. The phage network was built with these wGRR values, using the *networkx* and *graphviz* Phyton (v2.7) packages, and the *neato* algorithm.

### Generation of capsule mutant

Isogenic capsule mutants were generated by an in-frame knock-out deletion of gene *wza* (strain #26, #24, #56, #37) or gene *wcaJ* (strain #57 and #58) by allelic exchange. Upstream and downstream sequences of the *wza* or *wcaJ* gene (> 500pb) were amplified using Phusion Master Mix then joined by overlap PCR. All primers used are listed in Table S5. The PCR product was purified using the QIAquick Gel Extraction Kit after electrophoresis in agarose gel 1% and then cloned with the Zero Blunt^®^ TOPO^®^ PCR Cloning Kit (Invitrogen) into competent *E. coli* DH5α strain. KmR colonies were isolated and checked by PCR. A positive Zero Blunt^®^ TOPO^®^ plasmid was extracted using the QIAprep Spin Miniprep Kit, digested for 2 hours at 37°C with ApaI and SpeI restriction enzymes and ligated with T4 DNA ligase overnight to digested pKNG101 plasmid. The ligation was transformed into competent *E. coli* DH5α pir strain, and again into *E. coli* MFD λ–pir strain (Ferrieres et al 2010), which was used as a donor strain for conjugation into *Klebsiella spp*. Conjugations were performed for 24 hours at 37°. Single cross-over mutants (transconjugants) were selected on Streptomycin plates (200 µg/mL). Plasmid excision was performed after a 48h culture incubated at 25°C and double cross-over mutants were selected on LB without salt plus 5% sucrose at room temperature. To confirm the loss of the plasmid, colonies were tested for their sensitivity to streptomycin and mutants were confirmed by PCR across the deleted region and further verified by Illumina sequencing.

### Data availability

Raw data is available in https://datadryad.org/.

## Supporting information

Mat Sup

## ACKNOWLEDGEMENTS

We thank Pedro Oliveira for making available the profiles for the RMS systems, Marie Touchon for help in the analysis of CRISPR, and Amandine Perrin for help with building pan-genomes.

## SUPPLEMENTAL FIGURE LEGENDS

**Figure S1. Phylogenetic tree of 254 Klebsiella strains analysed in this study.** The tree was built using the protein sequences of the 1116 families of the core genome of *Klebsiella spp*. Red squares on the outer part of the tree indicate the presence (full) or absence (empty) of CRISPR-systems. The next two columns indicate the total number of prophages (brown) and the number of intact prophages (blue). Green circles indicate bootstrap values less than 99% (for clarity purposes all other nodes are supported by 100% bootstrap values and are not labelled). The size of the circle is proportional to the bootstrap value.

**Figure S2. Characteristics of *Klebsiella spp* prophages. A.** Correlation between the average number of prophages per genome per species with average genome size of the species. Each point represents the average of all strains of one species. P values correspond to Spearman’s correlation. **B.** GC content of “intact” prophages and host genomes. Each dot represents individual genomes. *** P < 0.001, Wilcoxon test. **C.** Distribution of the length of “intact” prophages.

**Figure S3. Average number of prophages in the 100 species with most genomes sequenced.** All complete genomes were downloaded November 2016 from NCBI RefSeq (ftp://ftp.ncbi.nih.gov/genomes/)), regrouped by species, and the 100 most sequenced species were selected and all of their genomes were analysed by PHASTER for phage detection. Digits indicate the number of analyzed genomes. Dashed line indicated average of each category for the dataset.

**Figure S4. Growth of all 35 strains used in this study at different concentrations of mitomycin C (MMC).** The ecological source of the isolates is also indicated.

**Figure S5. Excision of temperate phages from their bacterial host. A.** PCR specific to different phages encoded in different strains, detected in induced supernatants (+) but also in non-induced (-). Filtered and PEG-precipitated supernatants were treated with double-straned DNAse (Thermo Scientific) for 10 minutes at 37°, and then 20 minutes at 65°, for enzyme degradation. 1 ul was used as PCR matrix, and 5ul of PCR were loaded on an agarose gel. Numbers correspond to the host genomes, as displayed in Figure 3. The number in parenthesis identifies the prophage in the genome. **B.** Recircularisation proof for several temperate phages. Primers were designed to match prophage regions at the borders, but in opposite genomic directions. Presence of a PCR product indicated that the phage has excised and recircularized.

**Figure S6. Prophages can excise and infect other *Klebsiella* strains**. **A.** Growth of BJ1 strain after the addition of different concentrations of purified phage (shades of green) produced by strain #54 in the presence (full line) or absence (dashed line) of citrate (0.2%). **B.** Serial dilutions of lysates from different strains on an overlay of strain BJ1. Numbers indicate the dilution. **C.** PFU per ml produced by PEG-precipitation and filtered-supernatants lysates of strain #54 which were induced and not induced by MMC. Dashed line indicates the limit of detection of our assay. All experiments were performed in triplicate and error bars correspond to standard deviation of independent biological replicates.

**Figure S7. Number of independent replicates in which a given infection was observed**. The infection matrix was built using three independent lysates over three independent overlays. In very few cases did the three independently-generated lysates infect (independent replicates). The graph reports the number of infections that were observed in one, two or three independent replicates. The panels are separated as to show infections from lysates from a bacteria with a different capsule locus type than that of the targeted bacteria (Diff, first panel) or from the same capsular locus type (Same, second panel).

**Figure S8. Phage similarity does not confer resistance to superinfection. A.** Network of prophages (intact & questionable) of 35 *Klebsiella* strains as calculated by the wGRR (indicated in red along the connexions), with a cut-off of 0.5. Node colors represent different capsule locus types (as in Figure 4). **B.** Correlation between the wGRR of phage genomes and their respective phage repressors.

**Figure S9. Number of spacers and R-M systems targeting intact prophages genomes from other strains. A.** Spacer sequences of each CRISPR-Cas array were identified and blasted against the genomes of all intact prophages. The colours in the matrix reflect the number of different spacers in a strain (x-axis) that matches a prophage in a given strain (y-axis). **B.** The sequence motifs targeted by each R-M system present in one strain but absent in another were identified and blasted against the genomes of all intact prophages. The colours in the matrix reflect the number of restriction sites in a strain (x-axis) that matches a prophage hosted in another strain (y-axis).

**Figure S10. Number of CFU/mL per day, per strain and per condition.** Numbers in grey banner indicate the strain. The area under the curve between the populations grown in the presence or absence of citrate (LB vs LB + citrate, and MMC vs MMC + citrate) was not significantly different for all strains (T-test, P > 0.05).

## SUPPLEMENTAL TABLE LEGENDS

**Table S1. Colicins and microcins identified in the strains experimentally tested in this study**. In bold are identified those profiles with a match higher than 55%.

**Table S2. HMM profiles and sequences used to detect depolymerases.** HMM profiles associated with bacteriophage-encoded depolymerases from multiple bacterial species as well as the genetic sequences of five experimentally validated depolymerases.

**Table S3. Prophage proteins matching an HMM profile for capsule depolymerases.** HMM profiles are described in Table S2. The depolymerases detected in the strains used experimentally in this study are indicated in the column ‘Strain #’ that matches the numbers in figure 3 and 4.

**Table S4. BLASTP protein hits from “intact” prophages against sequences of experimentally-validated capsule depolymerases.** Hits were detected using *blastp*, and filtering by the e-value (maximum 1e-5), identity (40%) and coverage (40%).

**Table S5. Primers used in this study.**

**Table S6. pVOGS associated with phage repressors.**

**Dataset S1. Genomes used in this study, prophages and CRISPR arrays detected and spacer sequences.**

