## Supplementary material for "Modular prophage interactions driven by capsule serotype select for capsule loss under phage predation": Mat Sup

#### Table of Contents

SUPPLEMENTAL FIGURES 2

SUPPLEMENTAL TABLES 12

SUPPLEMENTAL REFERENCES 25

### SUPPLEMENTAL FIGURES

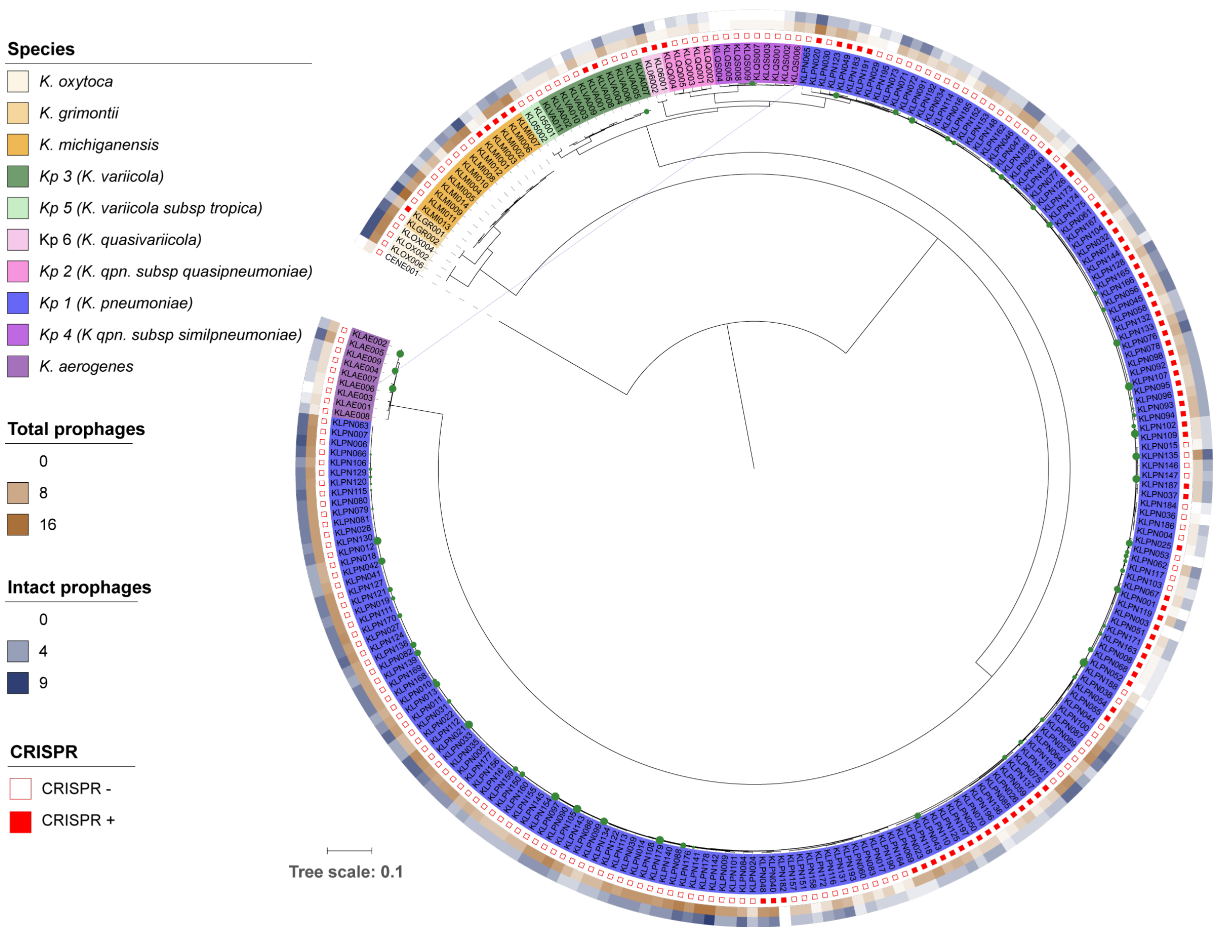

**Figure S1. Phylogenetic tree of 254 Klebsiella strains analysed in this study.** The tree was built using the protein sequences of the 1116 families of the core genome of *Klebsiella spp*. Red squares on the outer part of the tree indicate the presence (full) or absence (empty) of CRISPR-systems. The next two columns indicate the total number of prophages (brown) and the number of intact prophages (blue). Green circles indicate bootstrap values less than 99 (for clarity purposes). The size of the circle is proportional to the bootstrap value.

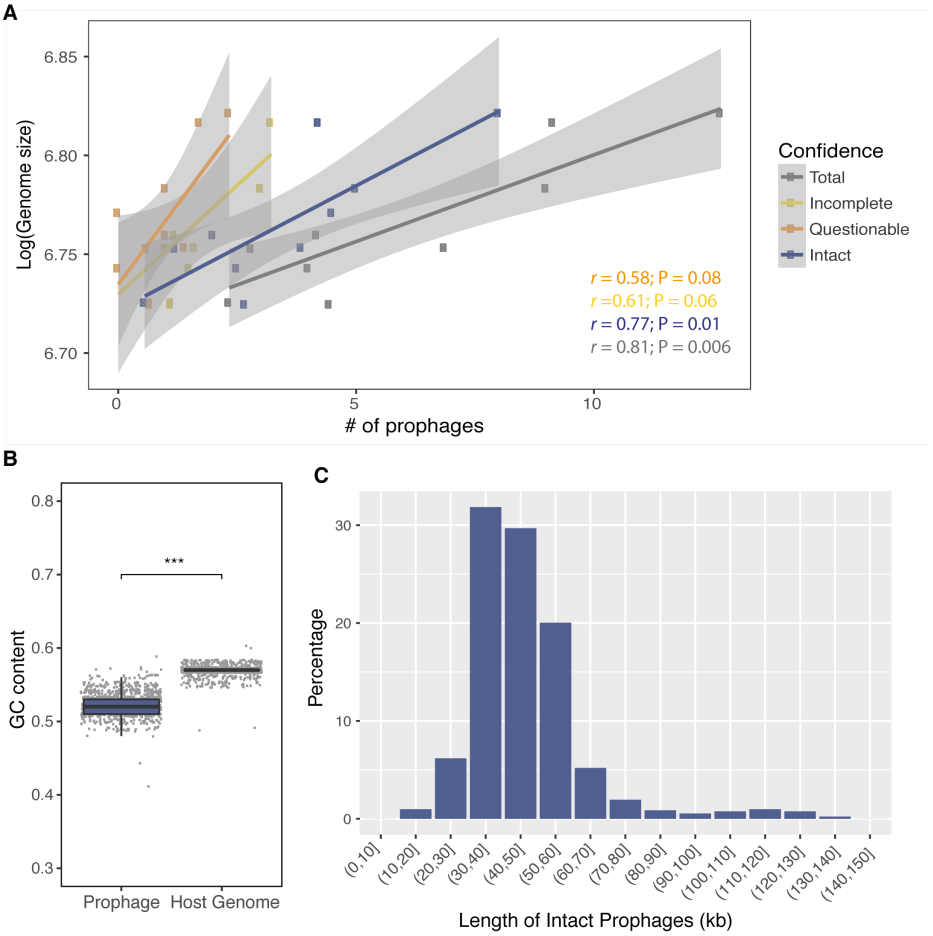

**Figure S2. Characteristics of *Klebsiella spp* prophages. A.** Correlation between the average number of prophages per genome per species with average genome size of the species. Each point represents the average of all strains of one species. P values correspond to Spearman's correlation. **B.** GC content of "intact" prophages and host genomes. Each dot represents individual genomes. *** P < 0.001, Wilcoxon test. **C.** Distribution of the length of "intact" prophages.

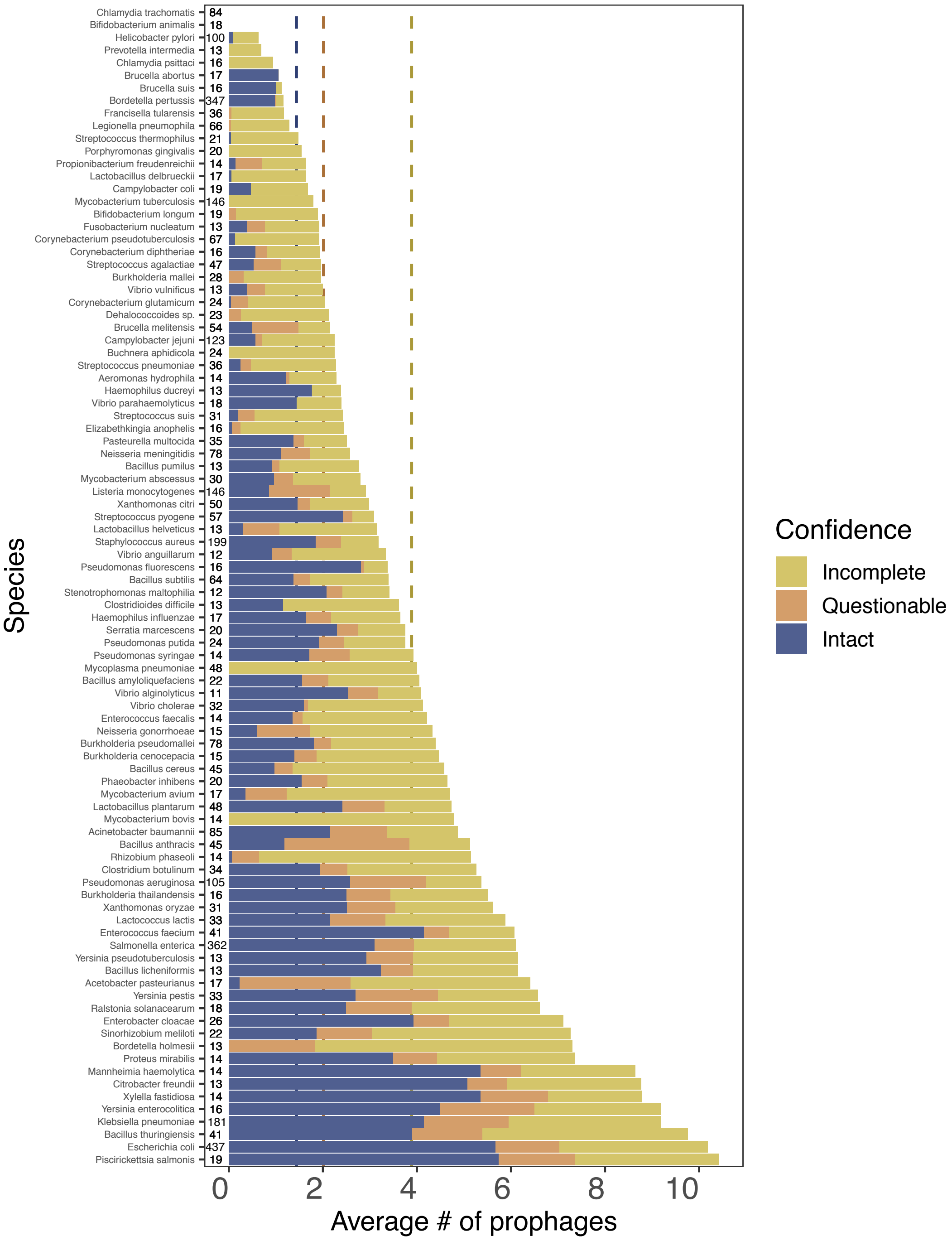

**Figure S3. Average number of prophages in the 100 species with most genomes sequenced.** All complete genomes were downloaded November 2016 from NCBI RefSeq (ftp://ftp.ncbi.nih.gov/genomes/)), regrouped by species, and the 100 most sequenced species were selected. Their genomes were analysed by PHASTER for phage detection. Digits indicate the number of analyzed genomes. Dashed line indicated average of each category for the dataset.

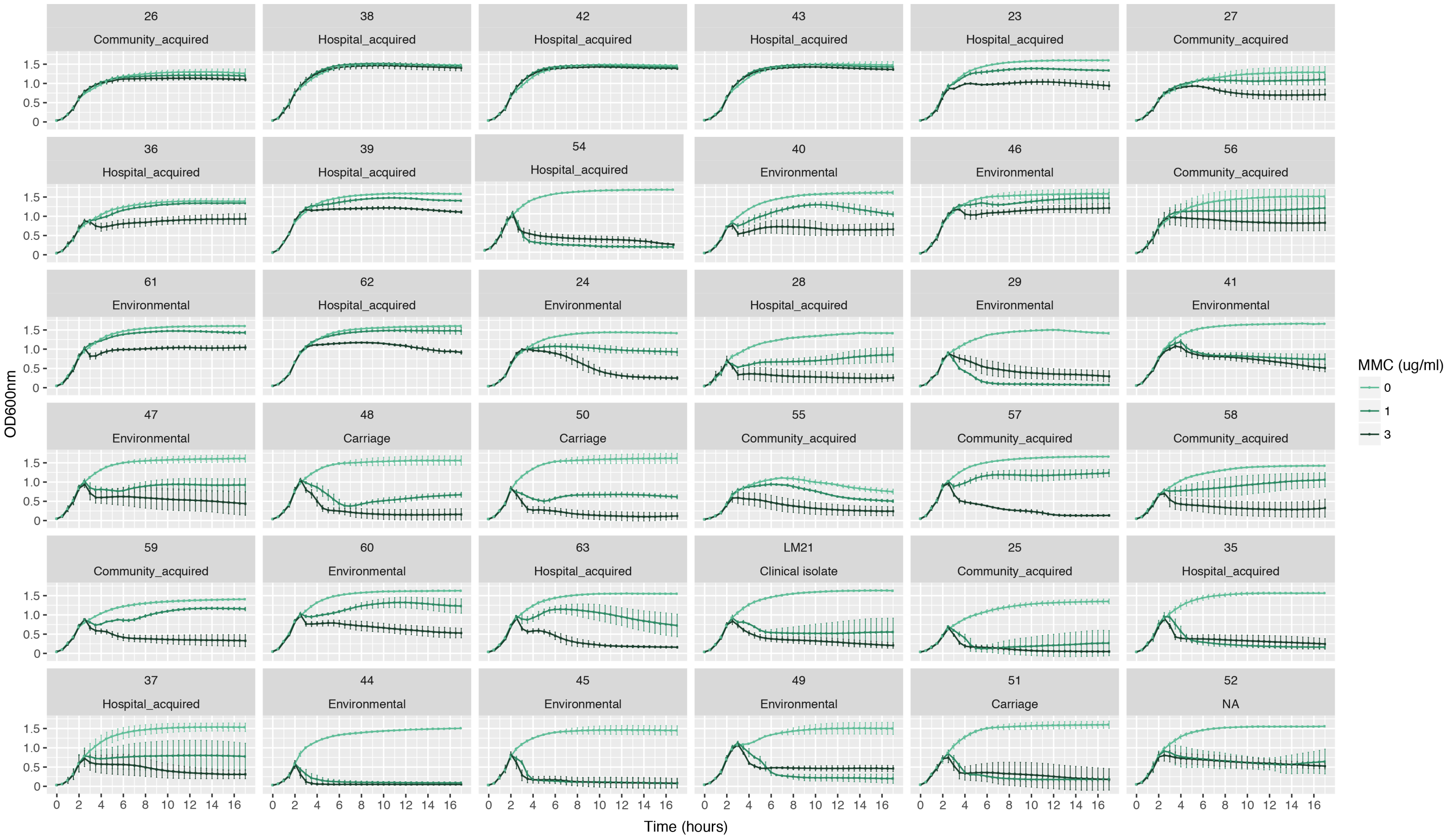

**Figure S4.** Growth of all 35 strains used in this study at different concentrations of mitomycin C (MMC). The ecological source of the isolates is also indicated.

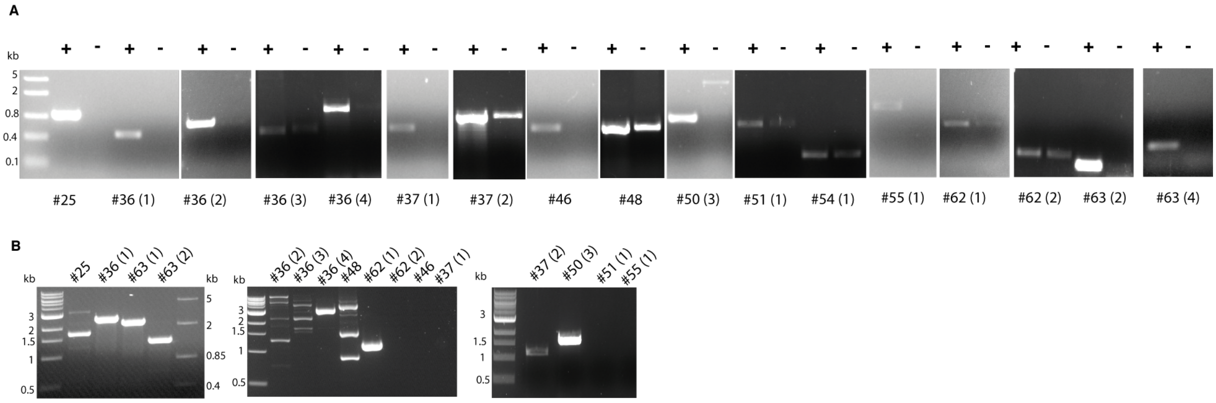

**Figure S5. Excision of temperate phages from their bacterial host. A.** PCR specific to different phages encoded in different strains, detected in induced supernatants (+) but also in non-induced (-). Filtered and PEG-precipitated supernatants were treated with double-stranded DNAse (Thermo Scientific) for 10 minutes at 37°, and then 20 minutes at 65°, for enzyme degradation. One $\mu$l was used as PCR matrix, and 5$\mu$l of PCR were loaded on an agarose gel. Numbers correspond to the host genomes, as displayed in Figure 3. The numbers in parenthesis identify the prophage in the genome. **B.** Recircularisation proof for several temperate phages. Primers were designed to match prophage regions at the borders, but in opposite genomic directions. Presence of a PCR product indicated that the phage has excised and recircularized.

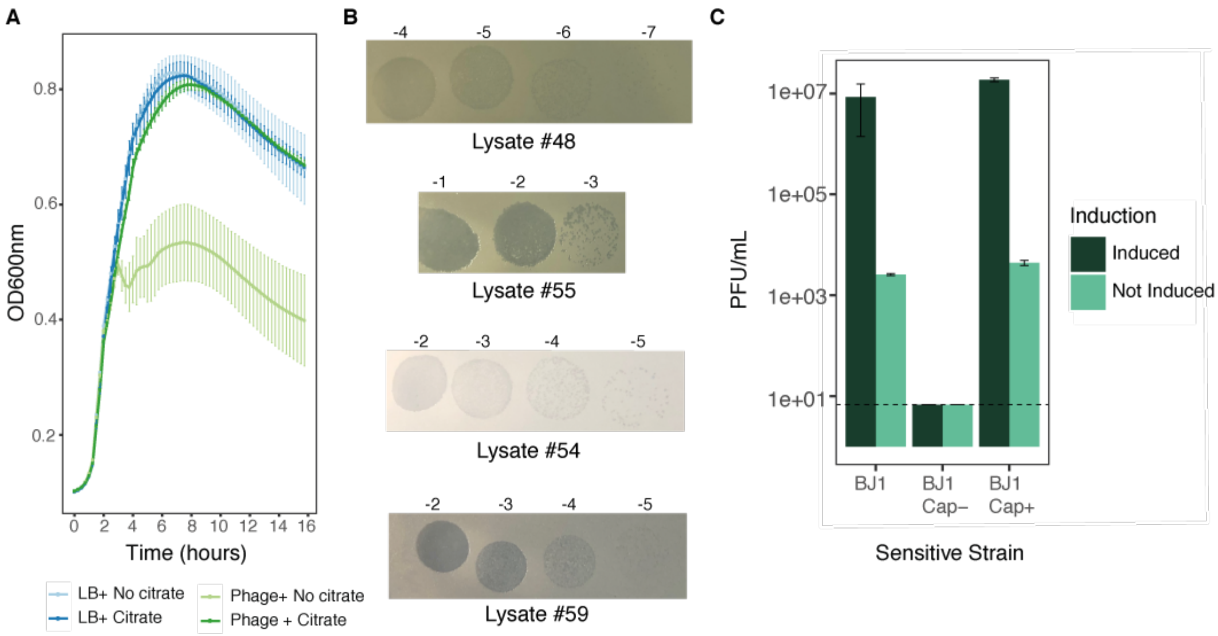

**Figure S6. Prophages can excise and infect other *Klebsiella* strains**. **A.** Growth of BJ1 strain after the addition of different concentrations of purified phage (shades of green) produced by strain #54 in the presence (full line) or absence (dashed line) of citrate (0.2%). **B.** Serial dilutions of lysates from different strains on an overlay of strain BJ1. Numbers indicate the dilution. **C.** PFU per ml produced by PEG-precipitation and filtered-supernatants lysates of strain #54 which were induced and not induced by MMC. Dashed line indicates the limit of detection of our assay. All experiments were performed in triplicate and error bars correspond to standard deviation of independent biological replicates.

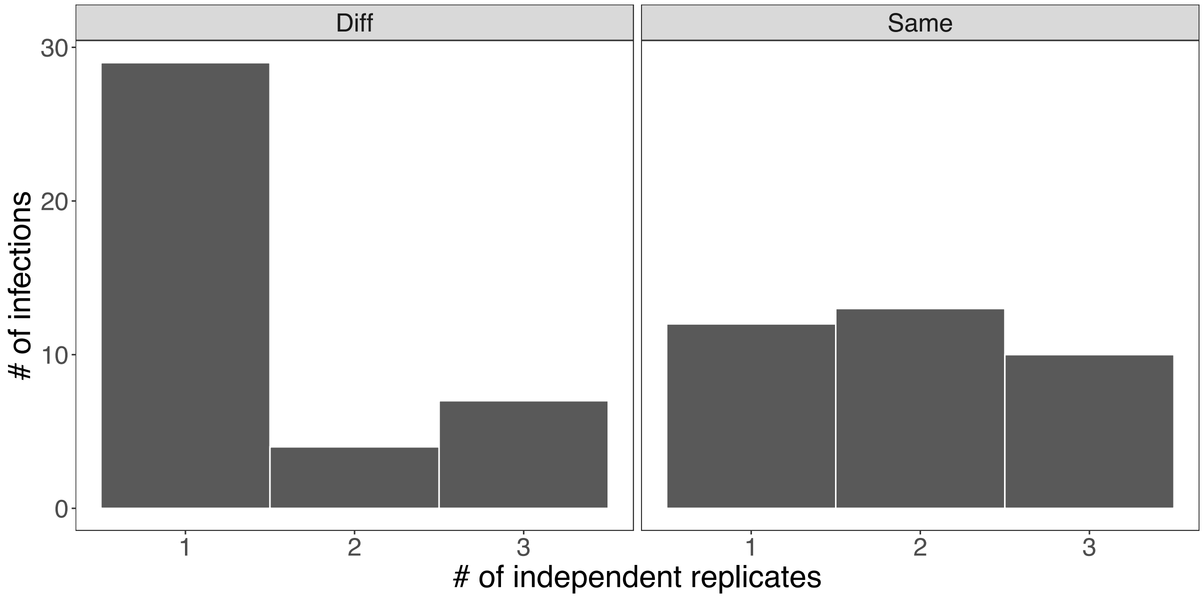

**Figure S7. Number of independent replicates in which a given infection was observed**. The infection matrix was built using three independent lysates over three independent overlays. In very few cases did the three independently-generated lysates infect (independent replicates). The graph reports the number of infections that were observed in one, two or three independent replicates. The panels are separated as to show infections from lysates from a bacteria with a different capsule locus type than that of the targeted bacteria (Diff, first panel) or from the same capsular locus type (Same, second panel).

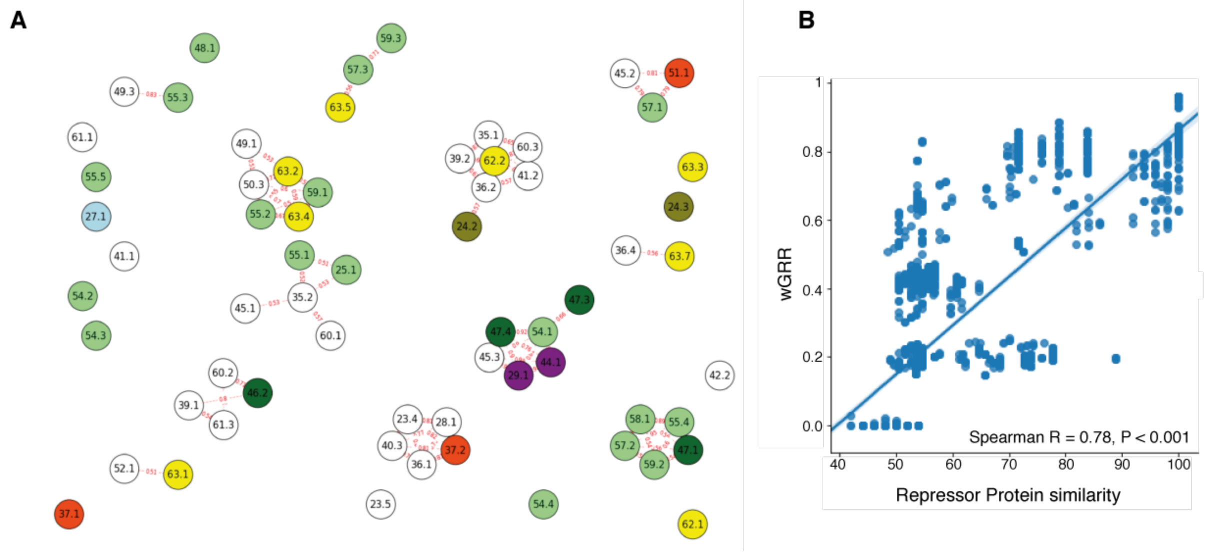

**Figure S8. Phage similarity does not confer resistance to superinfection. A.** Network of prophages (intact & questionable) of 35 *Klebsiella* strains as calculated by the wGRR (indicated in red along the connexions), with a cut-off of 0.5. Node colors represent different capsule locus types (as in Figure 4). **B.** Correlation between the wGRR of intact prophage genomes of the *Klebsiella* strains used in this study, and the protein sequence similarity of their respective phage repressors.

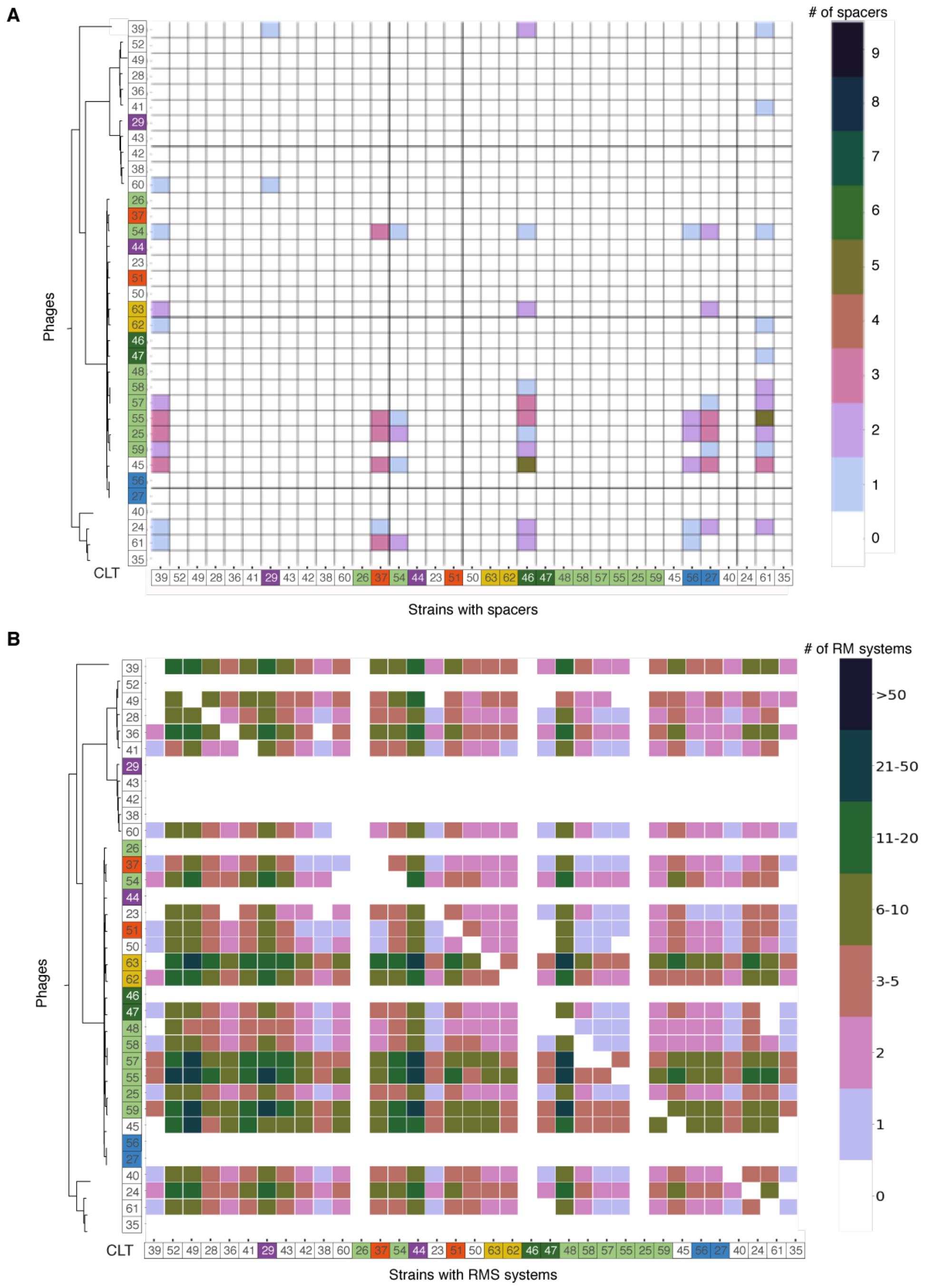

**Figure S9. Number of spacers and R-M systems targeting intact prophages genomes from other strains. A.** Spacer sequences of each CRISPR-Cas array were identified and blasted against the genomes of all intact prophages. The colours in the matrix reflect the number of different spacers in a strain (x-axis) that matches a prophage in a given strain (y-axis). **B.** The sequence motifs targeted by each R-M system present in one strain but absent in another were identified and blasted against the genomes of all intact prophages. The colours in the matrix reflect the number of R-M systems in a strain (x-axis) that matches a prophage hosted in another strain (y-axis).

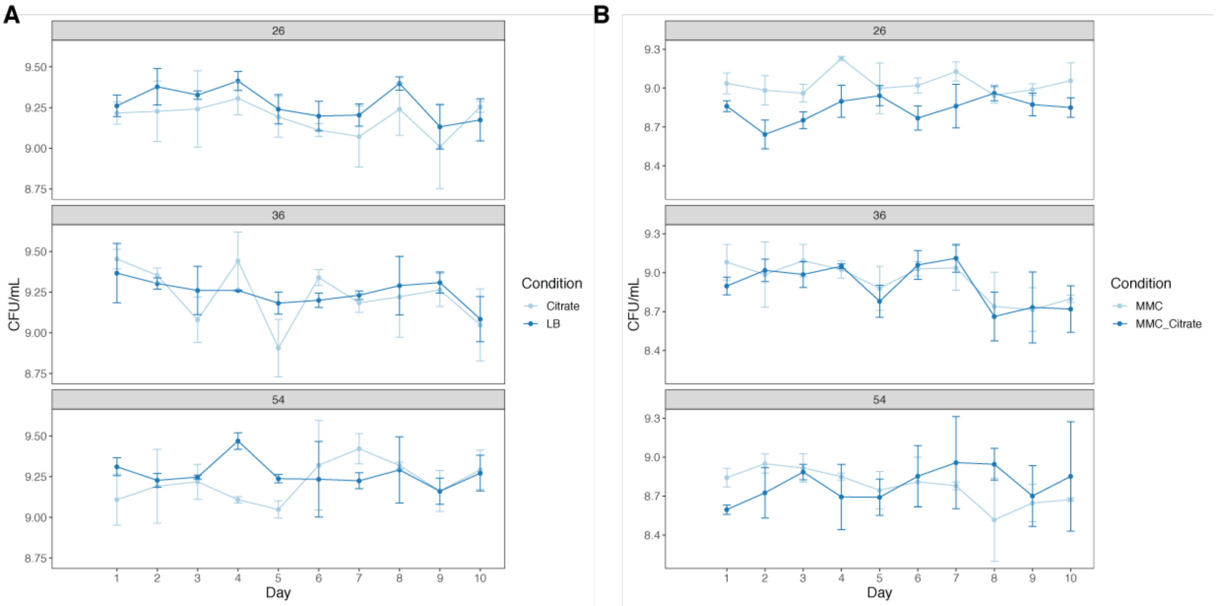

**Figure S10. Number of CFU/mL per day, per strain and per condition.** Numbers in grey banner indicate the strain. The area under the curve between the populations grown in the presence or absence of citrate (LB vs LB + citrate, and MMC vs MMC + citrate) was not significantly different for all strains (T-test, P > 0.05).

#### SUPPLEMENTAL TABLES

**Table S1. Colicins and microcins identified in the strains experimentally tested in this study. Indicated in bold are the profiles with a match higher than 55%.**

| Strain # | Contig # | Name | Start | End | Strand | Function | Match (%) | E-value | UniProt Profile | Strain infectivity |
|---|---|---|---|---|---|---|---|---|---|---|
| 63 | 4 | orf00041 | 15639 | 15788 | + | Lysis protein | **91.84** | 2.00E-25 | P02987 | Halo on 19 different strains |
| 63 | 4 | orf00042 | 15297 | 15554 | + | Cloacin immunity protein | **98.82** | 5.00E-55 | IMMC_ECOLX | |
| 63 | 4 | Colicin_E7 | 13602 | 15287 | + | Colicin_E7 | **71.02** | 1.00E-79 | Q47112 | |
| 23 | 2 | Colicin | 1465 | 3786 | + | Colicin | 48.85 | 9.00E-40 | | Mild halo on overlay of strain #47 |
| 23 | 2 | PyocinIm | 3788 | 4066 | + | Colicin-E9 immunity protein | 50.62 | 2.00E-22 | IMM9_ECOLX | |
| 23 | 3 | Colicin | 1465 | 3786 | + | Colicin | 48.85 | 9.00E-40 | | Mild halo on overlay of strain #47 |
| 23 | 3 | PyocinIm | 3788 | 4066 | + | Colicin-E9 immunity protein | 50.62 | 2.00E-22 | IMM9_ECOLX | |
| 48 | 59 | PyocinIm | 2499 | 2756 | + | Colicin-E9 immunity protein | 50 | 6.00E-23 | IMM9_ECOLX | Halo on 8 strains, 6 of which have the same CLT |
| 48 | 59 | Colicin | 2015 | 2497 | + | Colicin | 49.24 | 5.00E-43 | | |
| 44 | 127 | PyocinIm | 5265 | 5522 | + | Colicin-E9 immunity protein | 50.62 | 6.00E-23 | IMM9_ECOLX | No inhibition halo |
| 44 | 127 | orf00039 | 5093 | 5263 | + | Colicin-E7 | **55.36** | 1.00E-13 | Q47112 | |
| 44 | 127 | orf00042 | 4369 | 4980 | + | Colicin-D | 33.72 | 2.00E-14 | CEAD_ECOLX | |
| 27 | 281 | Microcin_M | 14588 | 14767 | - | Bacteriocin_Iic | 87.23 | 8.00E-27 | | No inhibition halo |
| 27 | 281 | LanT | 15041 | 17113 | - | Probable microcin-H47 secretion/processing ATP-binding protein mchF | 93.19 | 0 | MCHF_ECOLX | |
| 27 | 281 | HlyD | 17130 | 18191 | - | Microcin H47 secretion protein mchE | 92.07 | 0 | MCHE_ECOLX | |
| 27 | 281 | MicD | 18530 | 19021 | - | RTX-II toxin-activating lysine-acyltransferase ApxIIC | 34.57 | 9.00E-07 | P0A3I4 | |
| 27 | 281 | orf00052 | 21559 | 21828 | + | Bacteriocin_Iic | | | PF10439 | |
| 27 | 281 | Microcin_I47 | 22705 | 22872 | + | Microcin_I47 | 50 | 4.00E-11 | | |
| 27 | 281 | orf00067 | 25962 | 26249 | + | Microcin E492 immunity protein | **100** | 5.00E-61 | IM92_KLEPN | |
| 27 | 281 | Microcin_24 | 26329 | 26535 | + | Microcin_E492147.2 | **100** | 2.00E-41 | | |
| 27 | 281 | ABC | 34456 | 36174 | - | Putative multidrug export ATP-binding/permease protein | 33.4 | 3.00E-83 | P71082 | |

| | | | | | | | | | | |
|---|---|---|---|---|---|---|---|---|---|---|
| 27 | 281 | LanT | 36245 | 36469 | - | ABC-type bacteriocin/lantibiotic exporters, contain an N-terminal double-glycine peptidase domain | 33.4 | 4.00E-11 | Q2S9A9_HAHCH | |
| 36 | 2 | PyocinIm | 97234 | 97491 | + | Colicin-E9 immunity protein | 50.62 | 7.00E-23 | IMM9_ECOLX | Inhibition halo on #38 &#40. Lysogenization of #38 (Figure 4D) |
| 36 | 2 | Colicin | 95214 | 97232 | + | Colicin | 48.85 | 3.00E-40 | | |
| 28 | 76 | Colicin_E7 | 3933 | 5618 | + | Colicin_E7 | **71.02** | 1.00E-79 | Q47112 | Inhibition halo on #60, and mild halo on #59 and #61 |
| 28 | 76 | orf00023 | 5628 | 5885 | + | Cloacin immunity protein | **98.82** | 5.00E-55 | IMMC_ECOLX | |
| 28 | 76 | orf00024 | 5970 | 6119 | + | Lysis protein | **91.84** | 2.00E-25 | LYS0_ECOLX | |
| 52 | 89 | orf00019 | 1619 | 1774 | + | Lysis protein for colicin A | 60.78 | 8.00E-14 | LYS1_CITFR | |
| 52 | 89 | PyocinIm | 1254 | 1511 | + | Colicin-E9 immunity protein | 50 | 6.00E-23 | IMM9_ECOLX | |
| 52 | 89 | Colicin | 632 | 1252 | + | Colicin | 49.24 | 3.00E-42 | | |
| 42 | 25 | Colicin | 1474 | 3414 | + | Colicin | 48.85 | 2.00E-40 | | Mild halo on overlay of strain #47 |
| 42 | 25 | PyocinIm | 3416 | 3694 | + | Colicin-E9 immunity protein | 50.62 | 2.00E-22 | IMM9_ECOLX | |
| 43 | 40 | Colicin | 4914 | 5807 | + | Colicin | 49.24 | 2.00E-41 | | Mild halo on overlay of strain #47 |
| 43 | 40 | PyocinIm | 5809 | 6066 | + | Colicin-E9 immunity protein | | 6.00E-23 | IMM9_ECOLX | |
| 29 | 168 | PyocinIm | 3503 | 3760 | + | Colicin-E9 immunity protein | | 7.00E-23 | IMM9_ECOLX | Mild halo on overlay of strain #52, #60 and #62 |
| 29 | 168 | orf00024 | 2382 | 2564 | - | | | | | |
| 29 | 168 | Colicin | 2608 | 3501 | + | Colicin | 49.24 | 2.00E-41 | | |
| 39 | 3 | PyocinIm | 77904 | 78161 | + | Colicin-E9 immunity protein | 50.62 | 6.00E-23 | IMM9_ECOLX | Mild halo on overlay of strain #47 |
| 39 | 3 | orf00035 | 76783 | 76965 | - | | | | | |
| 39 | 3 | Colicin | 77009 | 77902 | + | Colicin | 49.24 | 2.00E-41 | | |
| 26 | 1 | Microcin_24 | 3768943 | 3769149 | - | Microcin_E492147.2 | **100** | 2.00E-41 | | Mild halo on overlay of strain #47 |
| 26 | 1 | orf00033 | 3769229 | 3769516 | - | Immunity Microcin | **100** | 5.00E-61 | IM92_KLEPN | |
| 26 | 1 | Microcin_I47 | 3772606 | 3772773 | - | MccI47_mchS21 | 49.2 | 3.00E-14 | | |
| 26 | 1 | orf00048 | 3773650 | 3773919 | - | Bacteriocin_Iic | | | | |
| 26 | 1 | MicD | 3776457 | 3776948 | + | RTX-II toxin-activating lysine-acyltransferase ApxIIC | 34.57 | 9.00E-07 | P0A3I4 | |
| 26 | 1 | HlyD | 3777575 | 3778348 | + | Microcin H47 secretion protein mchE | **93** | 8.00E-174 | MCHE_ECOLX | |
| 26 | 1 | LanT | 3778365 | 3780437 | + | Probable microcin-H47 secretion/processing ATP-binding protein mchF | **93.19** | 0 | MCHF_ECOLX | |
| 26 | 1 | Microcin_M | 3780711 | 3780890 | + | Bacteriocin_Iic | **87.23** | 8.00E-27 | | |

**Table S2. HMM profiles and sequences used to detect depolymerases.** HMM profiles associated with bacteriophage-encoded depolymerases from multiple bacterial species as well as the genetic sequences of five experimentally validated depolymerases.

| Enzyme class | Polymer | Profile | Predicted domains | Cut-offs |
|---|---|---|---|---|
| Sialidases | Sialidic acid | PF12218.6 | End_N_terminal | e-value maximum 1e-3, minimum profile coverage 30% |
| | | PF12217.6 | End_beta_propel | |
| | | PF12219.6 | End_tail_spike | |
| | | PF13884.4 | Peptidase_S74 | |
| | | PF11962.6 | Peptidase_G2 | |
| Levanases | Levan | PF00251.18 | Glyco_hydro_32N | |
| | | PF08244.10 | Glyco_hydro_32C | |
| Xylosidases | Xylan | PF01229.15 | Glyco_hydro_39 | |
| Dextranases | Dextran | PF13199.4 | Glyco_hydro_66 | |
| Peptidases | Poly-y-glutamate | PF05908.9 | DUF867 | |
| Hyaluronidases | Hyaluronate | PF07212.9 | Hyaluronidase_1 | |
| Pectin/pectate lysases | Galacturonate | PF00544.17 | Pec_lyase_C | |
| | | PF12708.5 | Pectate_lyase_3 | |
| Lipases | Triacylglycerols | PF14606.4 | Lipase_GDSL_3 | |

| Sequence | Description | Cut-offs | Reference |
|---|---|---|---|
| YP_009226010.1 | Tail fiber protein | e-value maximum 1e-5, minimum coverage 40%, minimum identity 40% | (Majkowska-Skrobek et al 2016) |
| BAP15746.1 | Hypothetical protein | | (Lin et al 2014) |
| BAQ02780.1 | Tail fiber | | (Pan et al 2015) |
| S1-1 | Tail spike | | (Pan et al 2017) |
| S2-6 | Tail spike | | |

**Table S3. Prophage proteins matching an HMM profile for capsule depolymerases.** HMM profiles are described in Table S2.

| Phage protein hit | Strain # | Host CLT | Profile | E-value best domain | Coverage (%) best domain |
|---|---|---|---|---|---|
| KLAE.0918.00009.4_00069 | | KL107 | Pectate_lyase_3 | 1.30E-13 | 77.21 |
| KLOX.0918.00009.8_00003 | | KL68 | Pectate_lyase_3 | 3.40E-11 | 85.58 |
| KLAE.0918.00005.3_00031 | | KL68 | Peptidase_S74 | 3.30E-11 | 98.28 |
| KLAE.0918.00007.7_00003 | | KL68 | Peptidase_S74 | 1.40E-09 | 93.10 |
| KLAE.0918.00009.3_00003 | | KL107 | Peptidase_S74 | 1.40E-09 | 93.10 |
| KLMI.0918.00001.2_00027 | | KL152 | Peptidase_S74 | 0.0002 | 91.38 |
| KLMI.0918.00007.3_00067 | | KL43 | Peptidase_S74 | 1.80E-07 | 91.38 |
| KLOX.0918.00004.11_00064 | | KL18 | Peptidase_S74 | 5.20E-08 | 96.55 |
| KLOX.0918.00005.11_00073 | | KL68 | Peptidase_S74 | 7.00E-11 | 96.55 |
| KLOX.0918.00005.8_00051 | | KL68 | Peptidase_S74 | 5.10E-10 | 93.10 |
| KLOX.0918.00006.13_00056 | | KL68 | Peptidase_S74 | 7.00E-11 | 96.55 |
| KLOX.0918.00006.8_00051 | | KL68 | Peptidase_S74 | 5.10E-10 | 93.10 |
| KLOX.0918.00007.12_00003 | | KL68 | Peptidase_S74 | 7.00E-11 | 96.55 |
| KLOX.0918.00007.5_00008 | | KL68 | Peptidase_S74 | 5.10E-10 | 93.10 |
| KLOX.0918.00008.11_00022 | | KL43 | Peptidase_S74 | 2.70E-10 | 96.55 |
| KLOX.0918.00009.13_00004 | | KL68 | Peptidase_S74 | 5.10E-10 | 93.10 |
| KLOX.0918.00010.10_00051 | | KL70 | Peptidase_S74 | 1.80E-07 | 91.38 |
| KLOX.0918.00010.2_00054 | | KL70 | Peptidase_S74 | 5.10E-10 | 93.10 |
| KLPN.0918.00002.2_00001 | | KL1 | Peptidase_S74 | 0.0002 | 91.38 |
| KLPN.0918.00005.2_00004 | | KL38 | Peptidase_S74 | 1.70E-07 | 94.83 |
| KLPN.0918.00009.3_00027 | | KL67 | Peptidase_S74 | 6.00E-05 | 93.10 |
| KLPN.0918.00009.5_00002 | | KL67 | Peptidase_S74 | 0.00012 | 93.10 |
| KLPN.0918.00012.2_00002 | | KL1 | Peptidase_S74 | 0.0002 | 91.38 |
| KLPN.0918.00017.3_00001 | | KL47 | Peptidase_S74 | 5.20E-07 | 96.55 |
| KLPN.0918.00018.5_00001 | | KL47 | Peptidase_S74 | 5.20E-07 | 96.55 |
| KLPN.0918.00020.1_00030 | | KL1 | Peptidase_S74 | 0.0002 | 91.38 |
| KLPN.0918.00028.1_00029 | | KL1 | Peptidase_S74 | 0.0002 | 91.38 |
| KLPN.0918.00032.1_00029 | | KL1 | Peptidase_S74 | 0.0002 | 91.38 |
| KLPN.0918.00033.2_00029 | | KL47 | Peptidase_S74 | 5.20E-07 | 96.55 |
| KLPN.0918.00036.3_00002 | | KL1 | Peptidase_S74 | 0.0002 | 91.38 |
| KLPN.0918.00038.3_00025 | #25 | KL2 | Peptidase_S74 | 4.80E-07 | 96.55 |
| KLPN.0918.00040.2_00002 | | KL30 | Peptidase_S74 | 4.50E-07 | 91.38 |
| KLPN.0918.00045.3_00025 | | KL2 | Peptidase_S74 | 4.80E-07 | 96.55 |
| KLPN.0918.00047.6_00017 | | KL17 | Peptidase_S74 | 5.20E-07 | 96.55 |
| KLPN.0918.00049.3_00001 | | KL2 | Peptidase_S74 | 4.30E-07 | 96.55 |
| KLPN.0918.00050.2_00001 | | KL30 | Peptidase_S74 | 2.10E-06 | 93.10 |
| KLPN.0918.00050.7_00070 | | KL30 | Peptidase_S74 | 3.90E-09 | 93.10 |

| | | | | | |
|---|---|---|---|---|---|
| KLPN.0918.00051.4_00027 | | KL30 | Peptidase_S74 | 2.10E-06 | 93.10 |
| KLPN.0918.00051.6_00097 | | KL30 | Peptidase_S74 | 3.90E-09 | 93.10 |
| KLPN.0918.00057.11_00066 | | KL19 | Peptidase_S74 | 1.80E-07 | 91.38 |
| KLPN.0918.00057.11_00071 | | KL19 | Peptidase_S74 | 7.00E-08 | 91.38 |
| KLPN.0918.00057.7_00020 | | KL19 | Peptidase_S74 | 3.10E-10 | 96.55 |
| KLPN.0918.00058.4_00002 | | KL21 | Peptidase_S74 | 0.00012 | 93.10 |
| KLPN.0918.00059.3_00002 | | KL14 | Peptidase_S74 | 0.00012 | 93.10 |
| KLPN.0918.00060.3_00002 | | KL14 | Peptidase_S74 | 0.00012 | 93.10 |
| KLPN.0918.00069.5_00003 | | KL107 | Peptidase_S74 | 1.70E-06 | 93.10 |
| KLPN.0918.00069.6_00002 | | KL107 | Peptidase_S74 | 0.0002 | 91.38 |
| KLPN.0918.00076.4_00012 | | KL27 | Peptidase_S74 | 1.80E-07 | 91.38 |
| KLPN.0918.00080.3_00051 | | KL64 | Peptidase_S74 | 3.90E-09 | 93.10 |
| KLPN.0918.00085.3_00030 | | KL2 | Peptidase_S74 | 3.90E-09 | 93.10 |
| KLPN.0918.00086.5_00080 | | KL64 | Peptidase_S74 | 3.90E-09 | 93.10 |
| KLPN.0918.00094.3_00002 | | KL1 | Peptidase_S74 | 0.0002 | 91.38 |
| KLPN.0918.00094.6_00003 | | KL1 | Peptidase_S74 | 3.90E-09 | 93.10 |
| KLPN.0918.00097.4_00079 | | KL2 | Peptidase_S74 | 3.90E-09 | 93.10 |
| KLPN.0918.00101.5_00021 | | KL64 | Peptidase_S74 | 3.70E-05 | 91.38 |
| KLPN.0918.00102.5_00067 | | KL64 | Peptidase_S74 | 0.0002 | 91.38 |
| KLPN.0918.00113.6_00053 | | KL51 | Peptidase_S74 | 3.90E-09 | 93.10 |
| KLPN.0918.00114.4_00017 | | KL64 | Peptidase_S74 | 3.90E-09 | 93.10 |
| KLPN.0918.00114.8_00083 | | KL64 | Peptidase_S74 | 6.20E-07 | 91.38 |
| KLPN.0918.00121.5_00011 | | KL64 | Peptidase_S74 | 1.70E-07 | 91.38 |
| KLPN.0918.00121.5_00017 | | KL64 | Peptidase_S74 | 6.30E-07 | 91.38 |
| KLPN.0918.00126.7_00008 | | KL64 | Peptidase_S74 | 0.0002 | 91.38 |
| KLPN.0918.00128.7_00075 | | KL105 | Peptidase_S74 | 5.50E-07 | 96.55 |
| KLPN.0918.00131.7_00001 | | KL64 | Peptidase_S74 | 5.20E-07 | 96.55 |
| KLPN.0918.00132.4_00002 | | KL21 | Peptidase_S74 | 0.00012 | 93.10 |
| KLPN.0918.00133.14_00039 | | KL47 | Peptidase_S74 | 5.20E-07 | 96.55 |
| KLPN.0918.00133.4_00039 | | KL47 | Peptidase_S74 | 5.20E-07 | 96.55 |
| KLPN.0918.00135.14_00039 | | KL47 | Peptidase_S74 | 5.20E-07 | 96.55 |
| KLPN.0918.00135.4_00039 | | KL47 | Peptidase_S74 | 5.20E-07 | 96.55 |
| KLPN.0918.00141.6_00003 | | KL64 | Peptidase_S74 | 1.80E-07 | 91.38 |
| KLPN.0918.00142.5_00012 | | KL64 | Peptidase_S74 | 6.30E-07 | 91.38 |
| KLPN.0918.00142.5_00017 | | KL64 | Peptidase_S74 | 6.30E-07 | 91.38 |
| KLPN.0918.00143.5_00056 | | KL64 | Peptidase_S74 | 6.30E-07 | 91.38 |
| KLPN.0918.00143.5_00062 | | KL64 | Peptidase_S74 | 4.60E-07 | 91.38 |
| KLPN.0918.00144.6_00080 | | KL64 | Peptidase_S74 | 6.30E-07 | 91.38 |
| KLPN.0918.00144.6_00085 | | KL64 | Peptidase_S74 | 6.30E-07 | 91.38 |
| KLPN.0918.00145.5_00087 | | KL64 | Peptidase_S74 | 6.30E-07 | 91.38 |
| KLPN.0918.00145.5_00094 | | KL64 | Peptidase_S74 | 6.30E-07 | 91.38 |
| KLPN.0918.00146.5_00053 | | KL64 | Peptidase_S74 | 6.30E-07 | 91.38 |

| | | | | | |
|---|---|---|---|---|---|
| KLPN.0918.00146.5_00058 | | KL64 | Peptidase_S74 | 7.40E-08 | 91.38 |
| KLPN.0918.00147.5_00023 | | KL64 | Peptidase_S74 | 6.30E-07 | 91.38 |
| KLPN.0918.00147.5_00028 | | KL64 | Peptidase_S74 | 6.30E-07 | 91.38 |
| KLPN.0918.00150.2_00001 | | KL30 | Peptidase_S74 | 4.80E-07 | 96.55 |
| KLPN.0918.00150.3_00096 | | KL30 | Peptidase_S74 | 3.90E-09 | 93.10 |
| KLPN.0918.00152.3_00062 | | KL10 | Peptidase_S74 | 0.00067 | 93.10 |
| KLPN.0918.00154.11_00048 | | KL107 | Peptidase_S74 | 3.90E-09 | 93.10 |
| KLPN.0918.00154.2_00033 | | KL107 | Peptidase_S74 | 5.20E-07 | 96.55 |
| KLPN.0918.00157.5_00061 | | KL64 | Peptidase_S74 | 3.90E-09 | 93.10 |
| KLPN.0918.00157.6_00097 | | KL64 | Peptidase_S74 | 6.30E-07 | 91.38 |
| KLPN.0918.00157.6_00103 | | KL64 | Peptidase_S74 | 4.50E-08 | 91.38 |
| KLPN.0918.00159.4_00002 | | KL21 | Peptidase_S74 | 0.00012 | 93.10 |
| KLPN.0918.00163.2_00002 | #62 | KL107 | Peptidase_S74 | 4.80E-07 | 96.55 |
| KLPN.0918.00164.8_00057 | | KL103 | Peptidase_S74 | 5.50E-07 | 96.55 |
| KLPN.0918.00165.2_00002 | | KL107 | Peptidase_S74 | 4.80E-07 | 96.55 |
| KLPN.0918.00166.6_00003 | | KL64 | Peptidase_S74 | 1.80E-07 | 91.38 |
| KLPN.0918.00167.7_00053 | | KL51 | Peptidase_S74 | 3.90E-09 | 93.10 |
| KLPN.0918.00168.5_00015 | | KL64 | Peptidase_S74 | 3.70E-07 | 91.38 |
| KLPN.0918.00168.5_00020 | | KL64 | Peptidase_S74 | 6.30E-07 | 91.38 |
| KLPN.0918.00181.2_00072 | | KL7 | Peptidase_S74 | 3.90E-09 | 93.10 |
| KLPN.0918.00183.5_00042 | | KL112 | Peptidase_S74 | 3.90E-09 | 93.10 |
| KLPN.0918.00189.4_00002 | | KL21 | Peptidase_S74 | 0.00012 | 93.10 |
| KLPN.0918.00191.4_00003 | #47 | KL124 | Peptidase_S74 | 3.90E-09 | 93.10 |
| KLPN.0918.00196.3_00005 | #45 | KL116 | Peptidase_S74 | 3.90E-09 | 93.10 |
| KLPN.0918.00197.1_00104 | #54 | KL2 | Peptidase_S74 | 3.90E-09 | 93.10 |
| KLQQ.0918.00001.2_00002 | #36 | KL35 | Peptidase_S74 | 4.00E-06 | 91.38 |
| KLQQ.0918.00002.2_00049 | #41 | KL125 | Peptidase_S74 | 6.10E-07 | 93.10 |
| KLQQ.0918.00005.3_00020 | #49 | KL138 | Peptidase_S74 | 5.40E-07 | 96.55 |
| KLQS.0918.00002.3_00002 | #60 | KL57 | Peptidase_S74 | 0.00012 | 93.10 |
| KLQS.0918.00005.1_00015 | #29 | KL111 | Peptidase_S74 | 5.60E-07 | 96.55 |
| KLQU.0918.00003.3_00001 | | KL153 | Peptidase_S74 | 5.20E-07 | 96.55 |
| KLVA.0918.00003.4_00002 | | KL16 | Peptidase_S74 | 7.70E-05 | 93.10 |
| KLVA.0918.00005.2_00002 | #24 | KL30 | Peptidase_S74 | 4.50E-07 | 91.38 |
| KLVA.0918.00006.4_00002 | | KL103 | Peptidase_S74 | 0.0001 | 93.10 |
| KLVA.0918.00006.7_00006 | | KL103 | Peptidase_S74 | 4.10E-06 | 91.38 |
| KLVA.0918.00007.2_00023 | | KL53 | Peptidase_S74 | 1.20E-06 | 93.10 |
| KLVA.0918.00008.2_00023 | | KL53 | Peptidase_S74 | 1.20E-06 | 93.10 |
| KLVA.0918.00009.2_00035 | | KL53 | Peptidase_S74 | 1.20E-06 | 93.10 |
| KLVA.0918.00010.1_00002 | #35 | KL31 | Peptidase_S74 | 0.00031 | 93.10 |
| KP06.0918.00001.2_00001 | #39 | KL51 | Peptidase_S74 | 5.20E-07 | 96.55 |

**Table S4. BLASTP protein hits from "intact" prophages against sequences of experimentally-validated capsule depolymerases.** Hits were detected using *blastp*, and filtering by the e-value (maximum 1e-5), identity (40%) and coverage (40%).

| Phage protein hit | Strain # | Host CLT | Profile | Identity (%) | Bitscore | Coverage (%) |
|---|---|---|---|---|---|---|
| KLOX.0918.00010.10_00052 | | KL70 | YP_009226010.1 | 40.88 | 714 | 74.70 |
| KLPN.0918.00168.5_00021 | | KL64 | YP_009226010.1 | 41.09 | 714 | 74.70 |
| KLPN.0918.00147.5_00029 | | KL64 | YP_009226010.1 | 41.09 | 714 | 74.70 |
| KLPN.0918.00146.5_00052 | | KL64 | YP_009226010.1 | 41.09 | 714 | 74.70 |
| KLPN.0918.00145.5_00086 | | KL64 | YP_009226010.1 | 41.09 | 714 | 74.70 |
| KLPN.0918.00144.6_00079 | | KL64 | YP_009226010.1 | 41.09 | 714 | 74.70 |
| KLPN.0918.00143.5_00055 | | KL64 | YP_009226010.1 | 41.09 | 714 | 74.70 |
| KLPN.0918.00142.5_00018 | | KL64 | YP_009226010.1 | 41.09 | 714 | 74.70 |
| KLPN.0918.00121.5_00018 | | KL64 | YP_009226010.1 | 41.09 | 714 | 74.70 |
| KLMI.0918.00007.3_00066 | | KL43 | YP_009226010.1 | 41.09 | 714 | 74.70 |
| KLPN.0918.00154.2_00032 | | KL107 | YP_009226010.1 | 40.88 | 712 | 74.70 |
| KLOX.0918.00010.2_00053 | | KL70 | YP_009226010.1 | 42.87 | 712 | 68.61 |
| KLPN.0918.00076.4_00013 | | KL27 | YP_009226010.1 | 40.99 | 712 | 74.70 |
| KLPN.0918.00034.2_00032 | | KL64 | YP_009226010.1 | 41.20 | 712 | 74.70 |
| KLPN.0918.00166.6_00004 | | KL64 | YP_009226010.1 | 40.99 | 711 | 74.70 |
| KLPN.0918.00141.6_00004 | | KL64 | YP_009226010.1 | 40.99 | 711 | 74.70 |
| KLPN.0918.00126.7_00007 | | KL64 | YP_009226010.1 | 40.99 | 711 | 74.70 |
| KLPN.0918.00102.5_00066 | | KL64 | YP_009226010.1 | 40.99 | 711 | 74.70 |
| KLPN.0918.00101.5_00024 | | KL64 | YP_009226010.1 | 40.99 | 711 | 74.70 |
| KLVA.0918.00009.2_00034 | | KL53 | YP_009226010.1 | 40.99 | 711 | 74.70 |
| KLVA.0918.00008.2_00022 | | KL53 | YP_009226010.1 | 40.99 | 711 | 74.70 |
| KLVA.0918.00007.2_00022 | | KL53 | YP_009226010.1 | 40.99 | 711 | 74.70 |
| KLVA.0918.00006.7_00007 | | KL103 | YP_009226010.1 | 40.80 | 711 | 74.70 |
| KLPN.0918.00150.2_00002 | | KL30 | YP_009226010.1 | 40.06 | 711 | 77.21 |
| KLQS.0918.00002.3_00003 | #60 | KL57 | YP_009226010.1 | 40.88 | 711 | 74.70 |
| KLPN.0918.00165.2_00003 | | KL107 | YP_009226010.1 | 40.77 | 711 | 74.70 |
| KLPN.0918.00163.2_00003 | #62 | KL107 | YP_009226010.1 | 40.77 | 711 | 74.70 |
| KLPN.0918.00152.3_00061 | | KL10 | YP_009226010.1 | 41.09 | 711 | 74.70 |
| KLPN.0918.00094.3_00003 | | KL1 | YP_009226010.1 | 40.88 | 711 | 74.70 |
| KLPN.0918.00057.11_00065 | | KL19 | YP_009226010.1 | 40.88 | 711 | 74.70 |
| KLPN.0918.00051.4_00026 | | KL30 | YP_009226010.1 | 40.99 | 711 | 74.70 |

| | | | | | | |
|---|---|---|---|---|---|---|
| KLPN.0918.00050.2_00002 | | KL30 | YP_009226010.1 | 40.99 | 711 | 74.70 |
| KLPN.0918.00036.3_00003 | | KL1 | YP_009226010.1 | 40.88 | 711 | 74.70 |
| KLPN.0918.00032.1_00028 | | KL1 | YP_009226010.1 | 40.88 | 711 | 74.70 |
| KLPN.0918.00020.1_00029 | | KL1 | YP_009226010.1 | 40.88 | 711 | 74.70 |
| KLPN.0918.00012.2_00003 | | KL1 | YP_009226010.1 | 40.88 | 711 | 74.70 |
| KLPN.0918.00002.2_00002 | | KL1 | YP_009226010.1 | 40.88 | 711 | 74.70 |
| KLMI.0918.00001.2_00026 | | KL152 | YP_009226010.1 | 40.88 | 711 | 74.70 |
| KLPN.0918.00114.8_00082 | | KL64 | YP_009226010.1 | 40.88 | 710 | 74.70 |
| KLVA.0918.00003.4_00003 | | KL16 | YP_009226010.1 | 40.88 | 709 | 74.70 |
| KLPN.0918.00157.6_00096 | | KL64 | YP_009226010.1 | 40.88 | 709 | 74.70 |
| KLPN.0918.00133.4_00038 | | KL47 | YP_009226010.1 | 40.77 | 709 | 74.70 |
| KLPN.0918.00133.14_00038 | | KL47 | YP_009226010.1 | 40.77 | 709 | 74.70 |
| KLPN.0918.00189.4_00003 | | KL21 | YP_009226010.1 | 40.67 | 708 | 74.70 |
| KLPN.0918.00159.4_00003 | | KL21 | YP_009226010.1 | 40.67 | 708 | 74.70 |
| KLPN.0918.00135.4_00038 | | KL47 | YP_009226010.1 | 40.77 | 708 | 74.70 |
| KLPN.0918.00135.14_00038 | | KL47 | YP_009226010.1 | 40.77 | 708 | 74.70 |
| KLPN.0918.00132.4_00003 | | KL21 | YP_009226010.1 | 40.67 | 708 | 74.70 |
| KLPN.0918.00131.7_00002 | | KL64 | YP_009226010.1 | 40.77 | 708 | 74.70 |
| KLPN.0918.00069.6_00003 | | KL107 | YP_009226010.1 | 40.77 | 708 | 74.70 |
| KLPN.0918.00058.4_00003 | | KL21 | YP_009226010.1 | 40.67 | 708 | 74.70 |
| KLPN.0918.00033.2_00028 | | KL47 | YP_009226010.1 | 40.77 | 708 | 74.70 |
| KLPN.0918.00018.5_00002 | | KL47 | YP_009226010.1 | 40.77 | 708 | 74.70 |
| KLPN.0918.00017.3_00002 | | KL47 | YP_009226010.1 | 40.77 | 708 | 74.70 |
| KLQQ.0918.00005.3_00019 | #49 | KL138 | YP_009226010.1 | 40.34 | 707 | 74.70 |
| KLPN.0918.00028.1_00028 | | KL1 | YP_009226010.1 | 40.77 | 707 | 74.70 |
| KLQQ.0918.00001.2_00003 | #36 | KL35 | YP_009226010.1 | 40.67 | 706 | 74.70 |
| KLPN.0918.00057.7_00021 | | KL19 | YP_009226010.1 | 40.45 | 706 | 74.70 |
| KLPN.0918.00009.3_00026 | | KL67 | YP_009226010.1 | 40.67 | 704 | 74.70 |
| KLOX.0918.00007.5_00009 | | KL68 | YP_009226010.1 | 42.59 | 704 | 68.13 |
| KLOX.0918.00006.8_00050 | | KL68 | YP_009226010.1 | 42.59 | 704 | 68.13 |
| KLOX.0918.00005.8_00050 | | KL68 | YP_009226010.1 | 42.59 | 704 | 68.13 |
| KLAE.0918.00009.3_00006 | | KL107 | YP_009226010.1 | 42.40 | 699 | 68.61 |
| KLAE.0918.00007.7_00006 | | KL68 | YP_009226010.1 | 42.40 | 699 | 68.61 |
| KLPN.0918.00069.5_00004 | | KL107 | YP_009226010.1 | 42.81 | 697 | 67.96 |

**Table S5. Primers used in this study.**

| Name | Direction | Use | Sequence |
|---|---|---|---|
| Wza mutant construction | | | |
| KL1.wza.600-5 | Forward | Construction of wza mutant in strain #56 | TAGCTTCTATGGGCAGATGG |
| KL1.wza.600-3 | Reverse | Construction of wza mutant in strain #56 | CTGCTTCATATACGGGTGGAGC |
| KL1.wza.atg-3 | Reverse | Construction of wza mutant in strain #56 | AATGTCACATCATTAGTAAACC |
| KL1.wza.out-5 | Forward | Verification of wza mutant in strain #56 | GTTTCACCTTCACGCCATTCC |
| KL1.wza.out-3 | Reverse | Verification of wza mutant in strain #56 | TTTACTGCCGTCATCACCACG |
| KL30.wza.600-5 | Forward | Construction of wza mutant in strain #24 | ATGAACCGGGTAACCAACTGG |
| KL30.wza.600-3 | Reverse | Construction of wza mutant in strain #24 | TATCTCTGGTGCTGATTGCTGG |
| KL30.wza.L5 | Forward | Construction of wza mutant in strain #24 | GGCTAACACAGCTGCTCAGGAATTGGCAA ATTTTGGTTTACTGATGATGTGACATTGAC ATATGTTTGACTCTGTATTAGTGG |
| KL30.wza.atg-3 | Reverse | Construction of wza mutant in strain #24 | ATGCACATCATCAGTAAACC |
| KL30.wza.out-5 | Forward | Verification of wza mutant in strain #24 | TTTCACCTTCACGCCGTTCC |
| KL30.wza.out-3 | Reverse | Verification of wza mutant in strain #24 | ACCTTTAATCTCACCGGCAC |
| KL2.wza.500-5 | Forward | Construction of wza mutant in strain #26 | TTAACTGGTATGTGGAAGCGCATG |
| KL2.wza.500-3 | Reverse | Construction of wza mutant in strain #26 | TCACCAACCATTCGACCAAAG |
| KL2.wza.5 | Forward | Construction of wza mutant in strain #26 | ATGTTTAGTACAATATTAATTGTTTGC |
| KL2.wza.L3 | Reverse | Construction of wza mutant in strain #26 | GCGAACGGCATATATTCCCTGTGCAAACA ATTAATATTGTACTAAACATAATGTCACA TCATCAGTAAATCAAAATTTGC |
| KL2.wza.out-5 | Forward | Verification of wza mutant in #26 | ACCGGGACAGATAACGAACC |
| KL2.wza.out-3 | Reverse | Verification of wza mutant in #26 | CACTTAACCTTGCCCATCCACG |
| WcaJ mutant construction | | | |

| | | | |
|---|---|---|---|
| KL2.wcaJ.L5 | Forward | Verification of wcaJ mutant in #57 and #58 | gttacgttaatgaaacttatcgacagatatgcttgtttataaatgagtgattgaattctagatgctccttaagacaagg |
| KL2.wcaJ.500-3 | Reverse | Verification of wcaJ mutant in #57 and #58 | CACCATACTCAATGCCGTTATGC |
| KL2.wcaJ.L3 | Reverse | Verification of wcaJ mutant in #57 and #58 | aattcaatcactcatttataaac |
| KL2.wcaJ.500-5 | Forward | Verification of wcaJ mutant in #57 and #58 | ttcttaaacttaagaacataagagc |
| KL2.wcaJ.out-5 | Forward | Verification of wcaJ mutant in #57 and #58 | GAATGGAATTGTTCTGCCTCTGAC |
| KL2.wcaJ.out-3 | Reverse | Verification of wcaJ mutant in #57 and #58 | CTCAGTTCACCTTCGTTCCATTCG |
| Phage detection & recircularization | | | |
| 54_Ph01-R_circ | Reverse | Tests for recircularisation of Prophage 1 in strain #54 | GGTGATCGCTTCCTGTGACGTGTTTTCC |
| 54_Ph01-F1_circ | Forward | Tests for recircularisation of Prophage 1 in strain #54 | GGCGTGTTCTGGATTTCTACC |
| 54_Ph01-F2_circ | Forward | Tests for recircularisation of Prophage 1 in strain #54. | TAATGAACCAGTGCCCGTCTCTGCCTTCC |
| 54_Ph03_F | Forward | Tests for recircularisation of Prophage 3 in strain #54 | CGAGATGGTACGCCGTTATG |
| 54_Ph03_R | Reverse | Tests for integrase, predicted 120bp (in genome) | GATTTCCTCCCTGTGTCGTATT |
| 54_Ph03_recirc | Forward | Tests for recircularisation of Prophage 3 in strain #54 | TTTCTCAAACAGCGGGATGG |
| 54_Ph01_R2 | Reverse | Tests for presence of Prophage 1 of strain #54. | GCCGATTTGTTGCGAGATCC |
| 54_Ph03_F2 | Forward | Tests for presence of Prophage 3 of strain #54. | GCGTCTGACAAACCGATAACAGC |
| 54_Ph03_R2 | Reverse | Tests for presence of Prophage 3 of strain #54. | ACGGCGTAAAGAAATGACAGTGCC |
| 54_Ph02_F | Forward | Tests for presence of Prophage 2 of strain #54. | AAGCCTACAAACTGACTGACG |
| 54_Ph02_R | Reverse | Tests for presence of Prophage 2 of strain #54. | TGACAAAGGAACAAGGTGAGG |
| 54_Ph02_Fcirc | Forward | Tests for recircularization of Prophage 2 of strain #54. | CGATACCTCTGACGCTCTGG |
| 54_Ph04_F | Forward | Tests for presence of Prophage 4 of strain #54. | GAAGGTAAGTTTGTCGGCCAGC |
| 54_Ph04_R | Reverse | Tests for presence of Prophage 4 of strain #54. | TCATCCGGCACATCTTCGAGG |
| 54_Ph04_Fcirc | Forward | Tests for recircularization of Prophage 4 of strain #54. | ATCGTTGTCAGTATCGGTGG |
| 54_Ph01_R3 | Reverse | Tests for recircularization of Prophage 3 of strain #54. | CTCAACGCCTGGGCTATTGC |
| 25_Ph01_F | Forward | Tests for presence (and recircularization) of Prophage 1 of strain #25. | GAATGCTGTATCCGGTGGTTGC |

| | | | |
|---|---|---|---|
| 25_Ph01_R | Reverse | Tests for presence of Prophage 1 of strain #25. | GTGCATGTTCGGTCAGGTGG |
| 25_Ph01_R_circ | Reverse | Tests for recircularisation of Prophage 1 of strain #25 | AACGGCTTCGTCGTACATCG |
| 36_Ph01_F | Forward | Tests for presence of Prophage 1 of strain #36. | CGGTAGTTCGATGCCTGTTCTTTCC |
| 36_Ph02_F | Forward | Tests for presence of Prophage 2 of strain #36. | CTGTACCACGGTCACCAAATCG |
| 36_Ph02_R | Reverse | Tests for presence of Prophage 2 of strain #36. | ACCGCCGATTACACAACACG |
| 36_Ph02_Fcirc | Forward | Tests for recircularization of Prophage 2 of strain #36. | GGCAACAACACGCGGATCTCC |
| 36_Ph03_F | Forward | Tests for presence of Prophage 3 of strain #36. | ATATCAGTGATCGACCGGGAGC |
| 36_Ph03_R | Reverse | Tests for presence of Prophage 3 of strain #36. | ATAAACTGTCTGATGGCGGTGG |
| 36_Ph03_Rcirc | Reverse | Tests for recircularization of Prophage 3 of strain #36. | GATGAGGGCTTTATTGTAGGTGG |
| 36_Ph04_F | Forward | Tests for presence of Prophage 4 of strain #36. | AAGGATTTCTCCTGCCACACC |
| 36_Ph04_R | Reverse | Tests for presence of Prophage 4 of strain #36. | GAATCCTTCTTTGCGCGTCG |
| 36_Ph04_Rcirc | Reverse | Tests for recircularization of Prophage 4 of strain #36. | GGGATACAAACCAACCTGACG |
| 36_Ph03_R2 | Reverse | Tests for recircularization of Prophage 3 of strain #36. | GTATACCCTGAAGTCTCGCTGG |
| 36_Ph04_R2 | Reverse | Tests for recircularization of Prophage 4 of strain #36. | GTGCGGTCAGCAAATACTGG |
| 37_Ph01_F | Forward | Tests for presence of Prophage 1 of strain #37. | AGGAAAGGATTTCTCTGAGAGC |
| 37_Ph01_R | Reverse | Tests for presence of Prophage 1 of strain #37. | GGCTATCCTGAACGAACTCAATCG |
| 37_Ph01_Fcirc | Forward | Tests for recircularization of Prophage 1 of strain #37. | ACCGATCTTCTCTACCCAGC |
| 37_Ph02_F | Forward | Tests for presence of Prophage 2 of strain #37. | CTAACGCATCCTGCAACTGATTTCC |
| 37_Ph02_R | Reverse | Tests for presence of Prophage 2 of strain #37. | GGCGGAATTGGTCTCGTTGC |
| 37_Ph02_Fcirc | Forward | Tests for recircularization of Prophage 2 of strain #37. | GCAAAGCCCAACCTGACACC |
| 37_Ph01_Fc2 | Forward | Tests for recircularization of Prophage 1 of strain #37. | ATCATGGAGGGTCAGCTGCTGG |
| 37_Ph02_Fc2 | Forward | Tests for recircularization of Prophage 2 of strain #37. | TGTTCATCATGGTTTGGGTGACG |
| 37_Ph02_refFc | Forward | Tests for recircularization of Prophage 2 of strain #37 | TAAAGCGTTCTCAGGTTTCCTCGGG |
| 37_Ph02_refRc | Reverse | Tests for recircularization of Prophage 2 of strain #37 | TGGCTGGTAACGTGATCTTAATGC |
| 46_Ph01_F | Forward | Tests for presence of Prophage of strain #46. | TGGGCTCATCAGCATTGAAGG |
| 46_Ph01_R | Reverse | Tests for presence of Prophage of strain #46. | CCATGTTCACGCCAGTGTCG |
| 46_Ph01_Fcirc | Forward | Tests for recircularization of Prophage of strain #46. | TTGCCAGAATGGCCTGATTTCG |

| | | | |
|---|---|---|---|
| 46_Ph01_Fc2 | Forward | Tests for recircularization of Prophage 1 of strain #46 | TACAGGAAGGATACGGATTTCAGC |
| 48_Ph01_F | Forward | Tests for presence of Prophage of strain #48. | ACAGGTGAGCGTAGACCATCG |
| 48_Ph01_R | Reverse | Tests for presence of Prophage of strain #48. | ATTCTCTGTTGGTTGCCCTGC |
| 48_Ph01_F_circ | Forward | Tests for recircularization of Prophage of strain #48. | CTCATACCTGGGTTGCTTGC |
| 48_Ph01_R_bis | Reverse | Tests for recircularization of Prophage 1 of strain #48 | TTCTGGAGCGGGAAGAGATCAGC |
| 50_Ph03_Fc | Forward | Tests for recircularization of Prophage 3 of strain #50 | CTTACTTTCGGGCCATGTTCAACG |
| 50_Ph03_R | Reverse | Tests for presence of Prophage 3 of strain #50 | CGGATTTACGACCAGCATCGAAC |
| 50_Ph03_F | Forward | Tests for presence of Prophage 3 of strain #50 | GATATGGCAGACCAGAGGTTACG |
| 51_Ph01_Fc | Forward | Tests for recircularization of Prophage 1 of strain #51 | GCGTACTGGCTGGATCTTAATTTCG |
| 51_Ph01_R | Reverse | Tests for presence of Prophage 1 of strain #51 | CGAGAAGCATACCCAGTTCAACC |
| 51_Ph01_F | Forward | Tests for presence of Prophage 1 of strain #51 | GCCTTACCAAATAACAACCCTCG |
| 55_Ph01_refFc | Forward | Tests for recircularization of Prophage 2 of strain #55 | GGATTTCTGCGGTTACGTTGTGC |
| 55_Ph01_refR | Reverse | Tests for presence of Prophage 1 of strain #55 | GCACTGTCATTTCTTTACGCCGTC |
| 55_Ph01_refF | Forward | Tests for presence of Prophage 1 of strain #55. | CTCTCCCATCCTCCCATTTCCTG |
| 62_Ph01_F | Forward | Tests for presence of Prophage 1 of strain #62. | AAGCCAGGGATGAACTGATAGC |
| 62_Ph01_R | Reverse | Tests for presence of Prophage 1 of strain #62. | ACAGCACACCGATTTCTTCC |
| 62_Ph01_Fcirc | Forward | Tests for recircularization of Prophage 1 of strain #62. | AGCAACAGCTCGCAAATCTCC |
| 62_Ph02_F | Forward | Tests for presence of Prophage 2 of strain #62. | ACGGTTAAACAGCGAGAAGC |
| 62_Ph02_R | Reverse | Tests for presence of Prophage 2 of strain #62. | CAGTATCACCAGTACCCAGC |
| 62_Ph02_F circ | Forward | Tests for recircularization of Prophage 2 of strain #62. | AGTCCTTTCCACTGCTTACC |
| 62_Ph02_Fcirc_bis | Forward | Tests for recircularization of Prophage 2 of strain #62. | TGCCAGTTTGTTTGCTTTCGTCC |

**Table S6. pVOGS associated with phage repressors.**

| **pVOG** |
|---|
| VOG0523 |
| VOG4990 |
| VOG4613 |
| VOG0126 |
| VOG4543 |
| VOG9234 |
| VOG0988 |
| VOG1681 |
| VOG2312 |
| VOG7724 |
| VOG0286 |
| VOG5346 |
| VOG9725 |
| VOG0513 |
| VOG8287 |
| VOG4597 |
| VOG6627 |
| VOG11077 |
| VOG8296 |
| VOG4696 |
| VOG4908 |
| VOG2167 |
| VOG9374 |
| VOG4614 |
| VOG5206 |
| VOG1658 |
| VOG0743 |
| VOG6362 |
